# Xenograft supervision and joint methylation-fragment-end encoding define a reusable molecular scoring layer for cell-free tumor DNA

**DOI:** 10.64898/2026.10.26.761533

**Authors:** Xuchao Zhang, Xiaojun Sun, Rui Wang, Ningning Zhao, Fang Liu, Xuhua Li, Jian Huang, Jiamin Sun, Han Li, Shang Lou, Haoyi Wang, Hailong Zhao, Han Shu, Peimeng Zhen, Hongcang Gu, Jiantao Shi, Tao Wang, Fan Zhang

## Abstract

Circulating tumor DNA is often scarce in plasma, and regional methylation summaries can obscure informative differences between individual molecules. Here we present ctDNA-BERT, a Transformer that scores individual bisulfite reads without predefined differentially methylated regions by jointly encoding CpG methylation state, nucleotide context and the fragment 3^***′***^ end. Xenografts provide molecule-level tumor supervision. The resulting frozen scorer supported tumor-fraction estimation in synthetic and patient-derived xenograft mixtures and enriched tumor-concordant methylation and chromatin-accessibility signals in independent hepatocellular carcinoma and stage III pan-cancer plasma without cohort-specific retraining. Features derived from selected reads discriminated stage I cancers from healthy controls in a retrospective cohort of 1,477 specimens, achieving an out-of-fold area under the receiver operating characteristic curve of 0.884. By separating molecular scoring from sample-level inference, ctDNA-BERT provides a reusable read-level scoring layer for cell-free DNA methylation analysis.

## 1 Introduction

Plasma cell-free DNA (cfDNA) carries genomic and epigenomic information from the cells that release it. DNA methylation is particularly informative because it reflects cell identity, enabling tissue mapping, multi-cancer detection and tissue-of-origin assignment [1–8]. The main limitation is abundance: circulating tumor DNA (ctDNA) is often a minor component of plasma, especially in early-stage disease [9], amid variable contributions from hematopoietic and other normal tissues [2, 10]. Accordingly, stage I sensitivity remains lower than that for advanced disease across current multi-cancer early-detection platforms [11, 12]. Cancer-associated methylation must therefore be detected before informative molecules are diluted by sample-level averaging.

Most cfDNA methylation workflows first restrict reads to differentially methylated regions (DMRs) or atlas-derived loci, then aggregate them into regional or sample-level summaries [1–3, 13–15]. This approach performs well when study samples resemble the discovery cohort and reference panel [16, 17], but imposes two constraints. Predefined regions exclude potentially informative tumor-derived reads outside the selected loci. Regional averaging also mixes rare tumor-derived molecules with abundant background reads and removes the physical linkage among CpG state, nucleotide context and fragment end. Once compressed to a regional mean, this molecule-level information cannot be recovered. The resulting gap is architectural rather than algorithmic: cfDNA analysis has no layer at which an individual molecule is assigned a tumor-derived score, so every downstream analysis inherits whichever summarization was applied upstream of it.

Individual reads retain this information. Methylation haplotypes and read-level epigenetic disorder capture signals that regional means can miss [18–20]. The same molecule also carries a fragment end shaped by chromatin organization and nuclease activity; end motifs, fragment size and coverage profiles provide independent information for cancer detection and tissue attribution [21–28]. Yet methylation and fragmentomic features are usually analyzed separately and combined only after sample-level summarization [29, 30].

Read-level models have begun to tackle this problem, but each addresses only part of it. DISMIR, MethylBERT and related methods established read-level scoring, but first restrict reads to predefined DMRs and so inherit the coverage and cohort-dependence limits of region selection [31–34]. Other Transformer models avoid DMR selection but couple read representations directly to end-to-end sample-level prediction, exposing no standalone molecular score that can be extracted, interpreted or reused [35–38]. Genomic language models show that self-supervised sequence pretraining can transfer across genomic tasks [39–44], but reference-sequence models encode neither CpG methylation state nor cfDNA fragment ends. What the field lacks, therefore, is not another read-level classifier but a molecular scoring layer: a component that assigns every eligible molecule a tumor-derived score, is fixed once and thereafter reused unchanged, so that molecular scoring and sample-level inference become separable design problems rather than a single end-to-end fit. Molecular enrichment has already been shown to improve ultrasensitive ctDNA monitoring [45], which is precisely the capability such a layer would make generally available. Two problems have kept this layer out of reach, and they are the problems this study sets out to solve.

The first problem is supervision. Plasma yields a label for the tube, not for the molecule, so read-level models have been trained either on labels inferred from the very regional statistics they were meant to supersede, or on bulk tumor tissue standing in for circulating tumor material. We obtain molecular labels by construction instead. In a xenograft, every sequenced fragment originates either from the engrafted human tumor or from the murine host, and the two are separated by their genomes rather than by any property the model will later learn: aligning each read against both hg19 and mm9 and retaining only unambiguous human assignments yields reads whose tumor origin is a fact of the experimental design rather than an output of a model. This turns the supervision bottleneck that the field has long acknowledged into an alignment problem with a definite answer, and it is what makes a molecule-level scorer trainable at all.

The second problem is representation. The CpG states of a molecule, its local nucleotide context and the position at which a nuclease cut it are physically linked on the same fragment, yet methylation and fragmentomic features are conventionally extracted by separate pipelines and reconciled only after each has been averaged to sample level, by which point the linkage no longer exists. We therefore write methylation into the token alphabet itself, so that a methylated CpG is a distinct lexical unit rather than an auxiliary channel appended to a sequence embedding, and supply the molecule’s 3^*′*^ terminal 6-mer as a paired segment carrying its own segment identity. Epigenetic state, sequence context and fragment end then interact through every self-attention operation of the encoder, rather than being combined after the information that relates them has been discarded.

These two solutions are realized in ctDNA-BERT, a DMR-independent Transformer that assigns every eligible bisulfite read a tumor-like score before any aggregation is performed. Methylated CpGs are encoded within the sequence alphabet, and the molecule s 3^*′*^ terminal 6-mer is provided as a paired segment so that methylation, sequence and fragment-end context interact throughout the encoder. Masked-language pretraining across 415 reduced-representation bisulfite libraries establishes a methylation-aware representation [46, 47]. Tissue fine-tuning separates tissue-derived reads from hematopoietic background, and xenograft fine-tuning supplies read-level tumor labels. The resulting frozen checkpoint supports tumor-fraction estimation, molecular enrichment and downstream feature construction without cohort-specific retraining. Scoring is deliberately decoupled from inference: the check-point is frozen once and never adapted again, so the applications reported below differ only in what is done with the scores afterward, never in the scorer that produced them. We used matched ablations and attention analyses to characterize the representation, benchmarked tumor-fraction recovery against four deconvolution methods and two read-level models, and tested transfer against pathologist-assigned tumor content in independent formalin-fixed specimens. We then applied the frozen scorer to independent hepatocellular carcinoma and pan-cancer plasma cohorts and evaluated selected-read features in a retrospective stage I case-control cohort of 1,477 specimens.

A high score denotes similarity to the tumor-derived training class, not definitive proof that an individual molecule is tumor-derived [11, 48].

## 2 Results

### 2.1 ctDNA-BERT encodes CpG state, nucleotide context and fragment end in one read-level representation

ctDNA-BERT assigns each eligible bisulfite read a tumor-like probability before regional or sample-level aggregation (**Fig. 1**). Because tumor, non-hematopoietic and hematopoietic cfDNA molecules are interleaved across the genome, the model treats the molecule rather than the locus as its unit of analysis (Fig. 1a). All cfDNA libraries in this study were generated using Primer-RRBS, which produces fragments bearing a shared MspI-defined 5^*′*^ terminus and the endogenous native breakpoint of the cfDNA molecule at the 3^*′*^ terminus (**Fig. 1b**). For each read, the strand-normalized methylation-aware body sequence and 6-nucleotide terminal sequence were supplied as paired segments. Converting both strands to a common orientation ensured that the same genomic context received one representation irrespective of the sequenced strand.

**Fig. 1.**
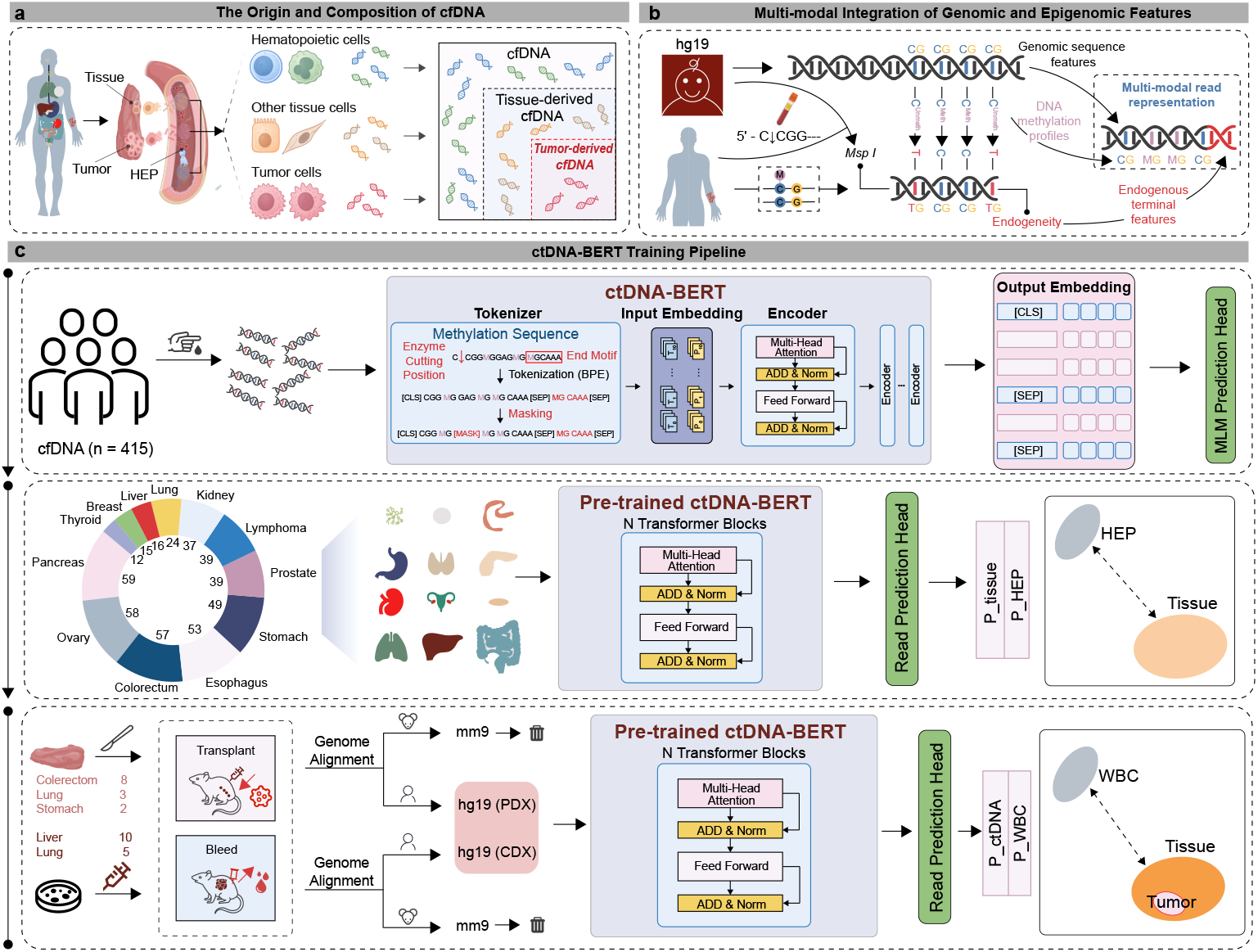
ctDNA-BERT integrates CpG state, nucleotide context and fragment end at read level. **a**, Plasma cfDNA comprises molecules released by tumor, non-hematopoietic tissue and hematopoietic cells; the model treats the molecule rather than the locus as its unit of analysis. **b**, Read representation. MspI digestion at C CGG defines the 5^*′*^ fragment terminus, while the 3^*′*^ terminus retains the endogenous native cfDNA breakpoint. The strand-normalized, methylation-aware body sequence and the 6-nucleotide terminal sequence are supplied as paired segments, with methylated CpGs encoded as MG or GM. **c**, Training pipeline. Top, methylation-aware strings are tokenized by SentencePiece byte-pair encoding into two segments ([CLS] body [SEP] terminal motif [SEP]) and processed by a 12-layer bidirectional Transformer trained by masked language modeling on 415 Primer-RRBS cfDNA libraries. Middle, tissue fine-tuning separates tissue reads (12 organ sites, positive class) from the 177-library hematopoietic reference panel (HEP, negative class). Bottom, xenograft fine-tuning aligns PDX and CDX reads against the hg19 and mm9 genomes; mouse-assigned reads are discarded, and the retained human-assigned tumor reads are contrasted with WBC reads to fit the final tumor-versus-background head. Parameters are frozen after this step. HEP, hematopoietic reference panel; WBC, white blood cell; PDX, patient-derived xenograft; CDX, cell line-derived xenograft; MLM, masked language modeling; BPE, byte-pair encoding.

Methylation was encoded within the token alphabet rather than appended as an auxiliary feature (**Fig. 1c, upper panel**). Methylated CpGs entered the tokenizer as MG or GM units, whereas unmethylated CpGs and non-CpG positions retained strand-normalized nucleotide identities. SentencePiece byte-pair encoding produced a vocabulary of 4,098 tokens: 4,096 learned subwords and two methylation-bearing units. This approach avoids the positional redundancy of overlapping fixed-length k-mers and is consistent with current DNA language models [39, 40, 49, 50]. Each input comprised [CLS], body tokens, [SEP], terminal-motif tokens and [SEP], with token-type embeddings distinguishing the two segments. Inputs were padded or truncated to 64 tokens, and padded positions were masked. The terminal segment therefore entered self-attention jointly with CpG state and local sequence.

The bidirectional Transformer comprised 12 layers, a hidden size of 768, 12 attention heads, a feed-forward width of 3,072 and dropout of 0.1. Masked-language pretraining used 415 Primer-RRBS cfDNA libraries for 10 epochs, with 15% of token positions selected for prediction. The library composition is shown in Supplementary Fig. 1a. The checkpoint at 460,000 optimization steps initialized subsequent fine-tuning. A linear two-class head applied to the final [CLS] representation produced the read-level tumor probability used downstream.

Two supervised fine-tuning stages then specialize the pretrained encoder. The first stage learns to separate tissue-derived reads from the hematopoietic background that dominates normal plasma, using reads from 458 study tissue libraries spanning 12 organ sites as the positive class and the 177-library hematopoietic reference panel (HEP) as the negative class (Fig. 1c, middle panel). Five of the 463 study tissue libraries, one from each of the breast, liver, lung, pancreas and thyroid groups, were reserved for testing at the tissue fine-tuning stage. The second stage supplies the molecular ground truth that plasma cannot provide. In a xenograft, a sequenced fragment has only two possible sources, the engrafted human tumor and the murine host, and these differ by genome rather than by epigenetic state; a read assigned unambiguously to the human genome is therefore tumor-derived by construction and not by inference. Critically, the label is independent of methylation, of fragment length and of every other feature the model is being trained to use, which is what makes it admissible as supervision for exactly those features. Reads from patient-derived (PDX) and cell line-derived (CDX) xenograft libraries were therefore aligned against both the human and mouse reference genomes [51]; reads assigned to the mouse genome were discarded, and the retained human-assigned tumor reads were contrasted with white-blood-cell (WBC) reads to fit the final tumor-versus-background head. This is the step that distinguishes ctDNA-BERT from read-level models supervised by labels derived from regional methylation statistics, which must assume that the regional summary is already correct in order to train the molecular model intended to supersede it. (**Fig. 1c, lower panel**). By design, the two stages use different negative classes — the hematopoietic reference panel in the first and white blood cells in the second. After xenograft fine-tuning, the neural read-level scorer was frozen for the plasma-enrichment analyses in Figs. 4–6. Matched ablations, the organ-system binary tasks and FFPE adaptation followed their separately specified training procedures; downstream sample-level classifiers were fitted within their training folds.

### 2.2 Methylation-aware encoding and the paired fragment-end segment contribute separably to read-level discrimination

Before benchmarking quantitative performance, we asked which components of the representation are responsible for separating tumor from background reads. We compared the complete configuration with three matched variants, each omitting exactly one element: masked-language pretraining, the paired terminal segment, or methylation-aware encoding. All four configurations followed identical training schedules and were evaluated on the same held-out labeled reads at 0, 4,000, 8,000, 12,000, 16,000 and 20,000 optimization steps. Score distributions for tumor and background reads are shown at each evaluation step in Fig. 2a. Read-level separation was additionally quantified by threshold-free ROC AUC using the same labeled evaluation reads. Early training-loss trajectories for the complete model and the three ablation configurations are shown in Supplementary Fig. 1b.

**Fig. 2.**
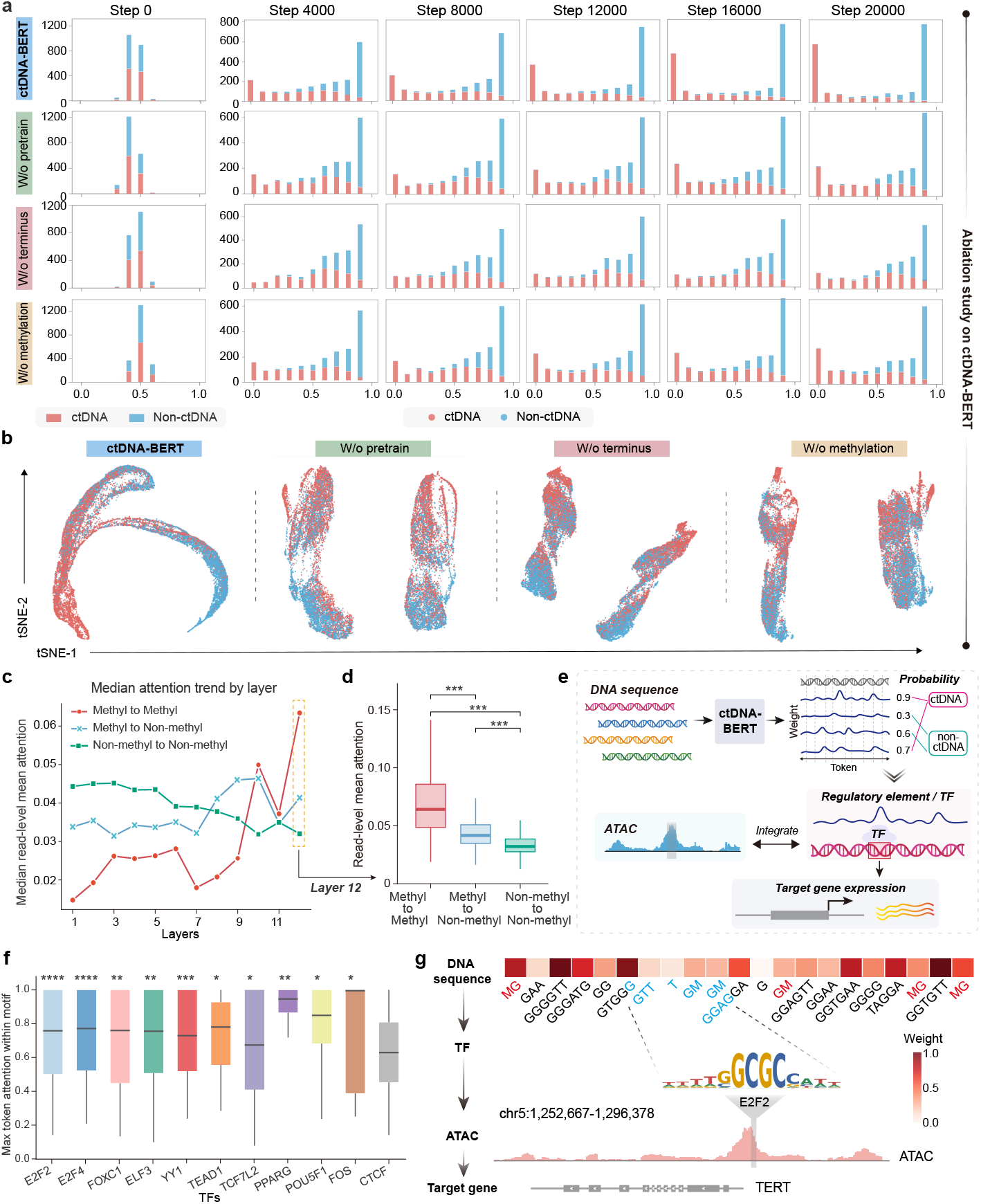
Matched ablations and attention analyses characterize the contribution of each input channel. **a**, Distributions of read-level model scores for tumor and background evaluation reads at six optimization steps (0, 4,000, 8,000, 12,000, 16,000 and 20,000), for the complete model and three matched ablations omitting pretraining, the paired terminal segment or methylation-aware encoding. All configurations used identical training schedules and the same held-out labeled reads. **b**, t-SNE projections of final-layer [CLS] representations, showing the clearest separation in the complete model. Adjusted Rand indices (ARI) were computed by k-means clustering (k = 2) in the full 768-dimensional space and compared with true labels. **c**, Median across reads of per-read mean attention (averaged over heads and eligible pairs) across 12 layers, for methylation-to-methylation, mixed and sequence-only token pairs. **d**, Layer-12 attention by pair category; paired two-sided tests with Benjamini-Hochberg adjustment. **e**, Workflow linking token attention to JASPAR transcription-factor motif intervals [52] and chromatin accessibility, with *CTCF* as the prespecified reference motif. **f**, Maximum within-motif token attention for *E2F2, E2F4, FOXC1, ELF3, YY1, TEAD1, TCF7L2, PPARG, POU5F1* and *FOS*, compared with the *CTCF* reference motif. Group sizes are *E2F2*, n = 428; *E2F4*, n = 285; *FOXC1*, n = 110; *ELF3*, n = 140; *YY1*, n = 91; *TEAD1*, n = 36; *TCF7L2*, n = 380; *PPARG*, n = 30; *POU5F1*, n = 24; *FOS*, n = 24; and *CTCF*, n = 502. These motifs were selected from 753 JASPAR motifs. Statistical comparisons used two-sided Mann–Whitney U tests with Benjamini–Hochberg adjustment. **g**, Single-read audit at the *TERT* promoter showing, for a representative 76-bp read, attention of 0.817 and 0.597 at two MG-bearing tokens and a mean of 0.381 (maximum, 0.940) across an *E2F2* motif, with the complete read lying within a called LIHC ATAC-seq open interval [53].

At 20,000 optimization steps, the complete model reached a read-level ROC AUC of 0.9753, compared with 0.8617 without pretraining, 0.9059 without the paired terminal segment and 0.9011 without methylation-aware encoding. The AUC gap associated with removal of pretraining increased from 0.0627 at 4,000 steps to 0.1136 at 20,000 steps. At 20,000 steps, removing the paired terminal segment and methylation-aware encoding reduced AUC by 0.0694 and 0.0742, respectively. The two channels therefore carry comparable and non-redundant weight, which is the operational meaning of encoding them jointly: neither is a refinement of the other, and a representation omitting either loses discrimination that the remaining channels do not recover. Final-layer [CLS] representations were visualized by t-SNE for descriptive comparison (Fig. 2b). We next examined attention across the 12 encoder layers. Token pairs were classified as methylation-to-methylation, mixed, or sequence-only. Attention was averaged across heads and over eligible pairs within each read before summarization, so that reads with more eligible pairs did not dominate (**Fig. 2c**). Methylation-to-methylation attention remained lowest through layer 9 and then rose sharply; at layer 12 it exceeded both mixed and sequence-only attention (0.062 versus 0.040 and 0.031; P < 0.001; Fig. 2d), consistent with the ablation results.

Attention also extended beyond CpG positions. Eligible LIHC reads were intersected with JASPAR transcription-factor motifs [52], with *CTCF* as the prespecified reference motif. Figure 2f shows maximum within-motif token attention for *E2F2, E2F4, FOXC1, ELF3, YY1, TEAD1, TCF7L2, PPARG, POU5F1* and *FOS*, with *CTCF* as the reference motif. Complementary comparisons for *LEF1* and *E2F1* are shown in Supplementary Fig. 1c. At the *TERT* promoter, a representative 76-bp read showed attention of 0.817 and 0.597 at two MG-bearing tokens, and a mean of 0.381 (maximum, 0.940) across an *E2F2* motif, with the complete read lying within a called LIHC ATAC-seq open interval ([53]; **Fig. 2g**). Additional token-to-token attention matrices and locus-level attention displays are shown in Supplementary Fig. 1d,e.

### 2.3 ctDNA-BERT recovers tumor fraction at read level across synthetic mixtures, held-out xenografts and pathology-matched FFPE specimens

We evaluated low-fraction tumor-read mixtures separately in WGBS and RRBS (Fig. 3a). The nominal target grid comprised 0.1–1.0% in increments of 0.1 percentage points and 1–10% in increments of 1 percentage point, with the shared 1% target counted once. WGBS comparisons included ctDNA-BERT, DISMIR, methAtlas and UXM; RRBS comparisons included ctDNA-BERT, UXM, methAtlas, cfSort and CelFEER. Upper panels summarize R^2^ and −ln(MSE), and lower panels show distributions of absolute estimation error over the available mixtures [16]. The complementary 10–90% analyses are shown in Supplementary Fig. 2a–d.

A second analysis examined low-fraction PDX mixtures for colorectal (COAD), lung (LUAD) and gastric (STAD) cancers across a 19-target grid spanning 0.1–10%, with MethylBERT and DISMIR included in the displayed comparison [31, 32]. Each box summarizes 19 target-specific MSE values, calculated across replicates at each target fraction. In this comparison, ctDNA-BERT had the lowest median target-specific MSE in colorectal (0.004735) and gastric mixtures (0.004731), compared with MethylBERT (0.010614 and 0.012097) and DISMIR (0.015791 and 0.014986), respectively. In lung mixtures, DISMIR had the lowest median MSE (0.000649), followed by ctDNA-BERT (0.004741) and MethylBERT (0.011295) (Fig. 3b). A complementary higher-fraction PDX analysis evaluated mixtures for the same three cancer types with a white-blood-cell background at nine target tumor fractions from 10% to 90%, in increments of 10 percentage points, with five replicates per tumor type and fraction. ctDNA-BERT estimates increased with the nominal tumor fraction across all three cancer types (Supplementary Fig. 2e).

**Fig. 3.**
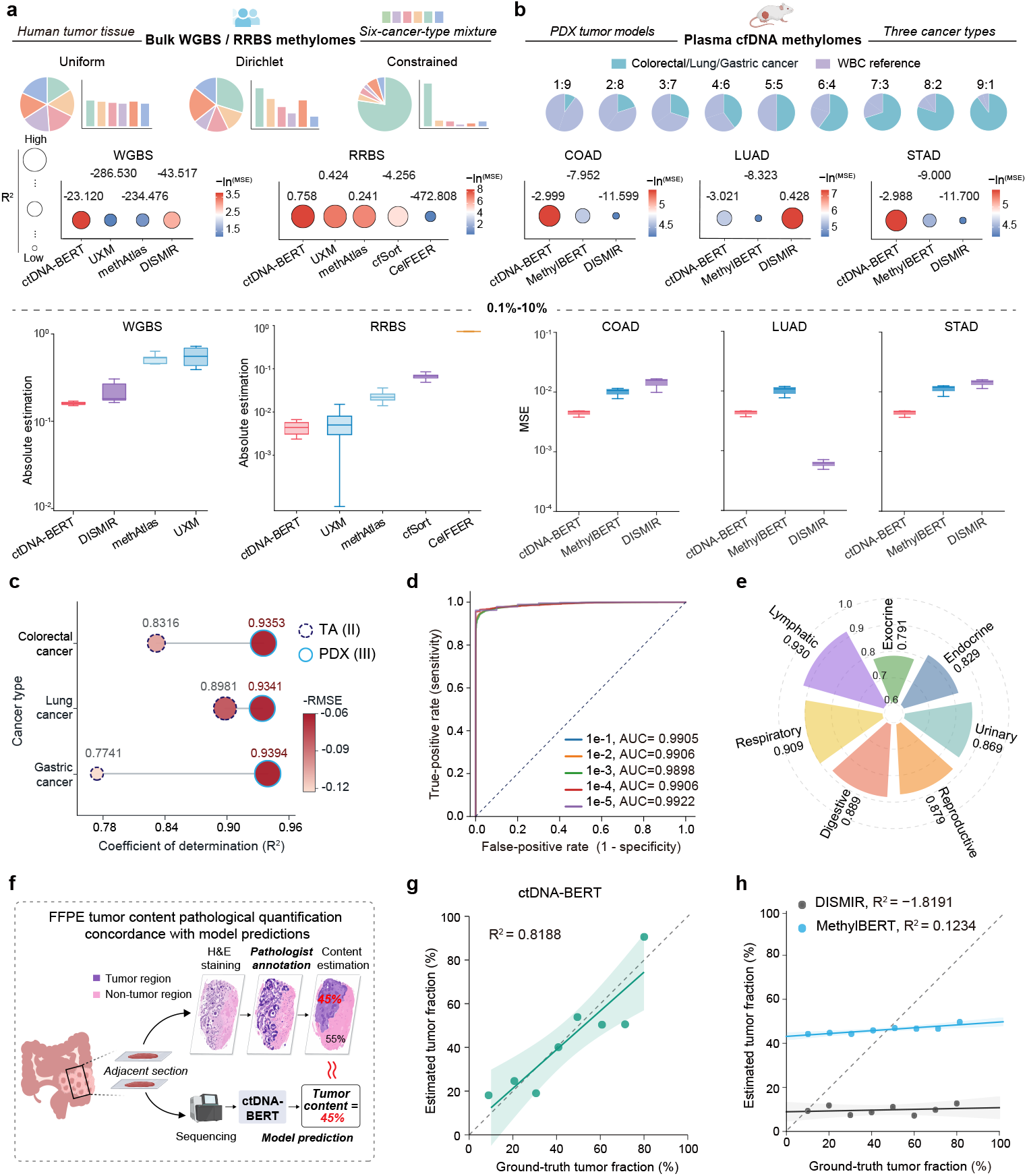
Tumor-fraction recovery in controlled mixtures, organ-system evaluations against the hematopoietic background, and independent liver FFPE tumor-content estimation. **a**, Platform-specific comparisons of tumor-fraction estimation in WGBS and RRBS mixtures. Upper panels summarize performance using R^2^ and color-coded −ln(MSE). Lower panels present the low-fraction comparisons across 19 distinct target fractions: 0.1–1.0% in increments of 0.1 percentage points and 1–10% in increments of 1 percentage point, with the shared 1% target counted once. Complementary higher-fraction analyses are shown in Supplementary Fig. 2a–d. Mixture structures: Uniform (equal source weights), Dirichlet (*α* = 0.05) and Constrained (one dominant source; realized weight, 0.621-0.740). ctDNA-BERT, methAtlas, UXM, cfSort and CelFEER analyzed identical files, with source-matched references where required. Top, estimated versus true tumor fraction; bottom, mean absolute error by method and mixture structure. **b**, Internal PDX-record mixture analysis for COAD, LUAD and STAD. Upper panels show R^2^, with color encoding −ln(MSE), over the 19-target 0.1– 10% grid. Lower panels show distributions of target-specific MSE values. MSE is calculated across five replicates per tumor type at every target fraction throughout the 0.1–10% grid, with the shared 1% target counted once; each box summarizes 19 target-specific MSE values. **c**, Contribution of xenograft fine-tuning, isolated by scoring the PDX mixtures with the tissue-fine-tuned checkpoint versus the xenograft-fine-tuned checkpoint. R^2^ was computed across fractions and replicates; color encodes −RMSE. **d**, Read-level ROC curves at tumor-read fractions of 10^*−*1^ to 10^*−*5^; AUC stayed at ∼0.99 across fractions (0.9898-0.9922). ROC AUC is prevalence invariant and does not establish a sample-level limit of detection. **e**, Seven independent organ-system-versus-HEP binary evaluations (same pretraining checkpoint and 177-library HEP reference), not a single seven-class classifier. Accuracies ranged from 0.791 (exocrine) to 0.930 (lymphatic). Lymphatic, n = 5; all other systems, n = 10. **f**, FFPE data-source workflow using adjacent sections for H&E staining and pathologist annotation, and corresponding material for sequencing and model-based tumor-content estimation. **g**, ctDNA-BERT tumor-content estimates for eight independent liver FFPE specimens after FFPE-specific fine-tuning and calibration on 23 non-liver specimens (8 colorectal, 8 gastric and 7 lung). Each point represents one specimen; the displayed agreement with the pathology reference is R^2^ = 0.8188. **h**, MethylBERT and DISMIR evaluated on the same eight liver FFPE specimens after their specified development procedures on the 23 non-liver specimens, with displayed R^2^ = 0.1234 and −1.8191, respectively. Dashed diagonals denote equality between reference and estimate. HEP, hematopoietic reference panel; PDX, patient-derived xenograft; RMSE, root-mean-square error.

Scoring the same mixtures with the tissue-fine-tuned checkpoint isolated the contribution of xenograft fine-tuning (**Fig. 3c**). Xenograft fine-tuning increased R^2^ from 0.8316 to 0.9353 in colorectal cancer, from 0.8981 to 0.9341 in lung cancer and from 0.7741 to 0.9394 in gastric cancer, with corresponding reductions in RMSE. Because the two checkpoints are identical except for whether molecule-level xenograft labels were used, this comparison isolates the value of the supervision itself rather than of any change in architecture, data volume or training schedule. Read-level discrimination remained stable as tumor-read prevalence decreased. ROC AUC was 0.9905, 0.9906, 0.9898, 0.9906 and 0.9922 as the labeled tumor-read fraction fell from 10^*−*1^ to 10^*−*5^ (**Fig. 3d**). This stability is expected, because ROC AUC is invariant to class prevalence [54].

Seven independent organ-system-versus-HEP evaluations tested tissue-background separation. Accuracies ranged from 0.791 to 0.930: lymphatic, 0.930; respiratory, 0.909; digestive, 0.889; reproductive, 0.879; urinary, 0.869; endocrine, 0.829; and exocrine, 0.791 (**Fig. 3e**). Each task used the same pretraining checkpoint and 177library HEP reference. These were independent binary tasks rather than a single seven-class classifier; the lymphatic estimate was based on only five specimens and is therefore imprecise.

Pathology-referenced tumor content was obtained from matched adjacent FFPE sections: an H&E-stained section provided pathologist annotation, while the corresponding section supplied DNA for sequencing and model-based estimation (Fig. 3f). An FFPE-adapted model was fine-tuned on pathologist-delineated tumor and adjacent non-tumor regions from 23 non-liver cancers (8 colorectal, 8 gastric and 7 lung) and calibrated to their pathologist-assigned tumor content [55]. After FFPE-specific fine-tuning and calibration, the model parameters and calibration mapping were frozen and applied to eight independent liver FFPE specimens (Fig. 3g). MethylBERT and DISMIR were trained, calibrated and evaluated on the same specimens. ctDNA-BERT achieved R^2^ = 0.8188 against pathologist-assigned tumor content (Fig. 3g), compared with 0.1234 for MethylBERT and −1.8191 for DISMIR, whose results are shown together in Fig. 3h. These are estimates of tumor content in fixed tissue, rather than calibrated plasma ctDNA fractions.

### 2.4 An uncalibrated frozen scorer enriches independent HCC plasma reads with convergent tumor-derived molecular signatures

We next asked whether the selected reads carried molecular features expected of tumor-derived material. The frozen scorer was applied without recalibration to plasma from 10 healthy controls and 76 patients with HCC, none of which were used for model training. Within each sample, the top 5% of reads formed the selected pool and the remainder the background pool (**Fig. 4a**). Terminal motifs, chromatin accessibility, tissue-level methylation and promoter methylation were then evaluated in both pools; none of these annotations had been available during training.

Fragment-end composition differed between pools. *CCAG* motifs were enriched among selected reads, whereas *TTTT* and *TTTA* were enriched among background reads (**Fig. 4b**), consistent with reported cancer-associated end-motif shifts [24, 27].

Selected reads also overlapped LIHC ATAC-seq peaks at a median density of 0.90 per 10^4^ valid reads per megabase of peak territory, compared with 0.40 for background reads (2.25-fold; P = 0.0002; [53]; Fig. 4c). Using an LIHC peak subset chosen for stage-associated behavior, an exploratory analysis showed that median chromatin overlap increased from healthy controls through stage IV HCC (n = 10, 43, 14, 13 and 6; **Fig. 4d**).

**Fig. 4.**
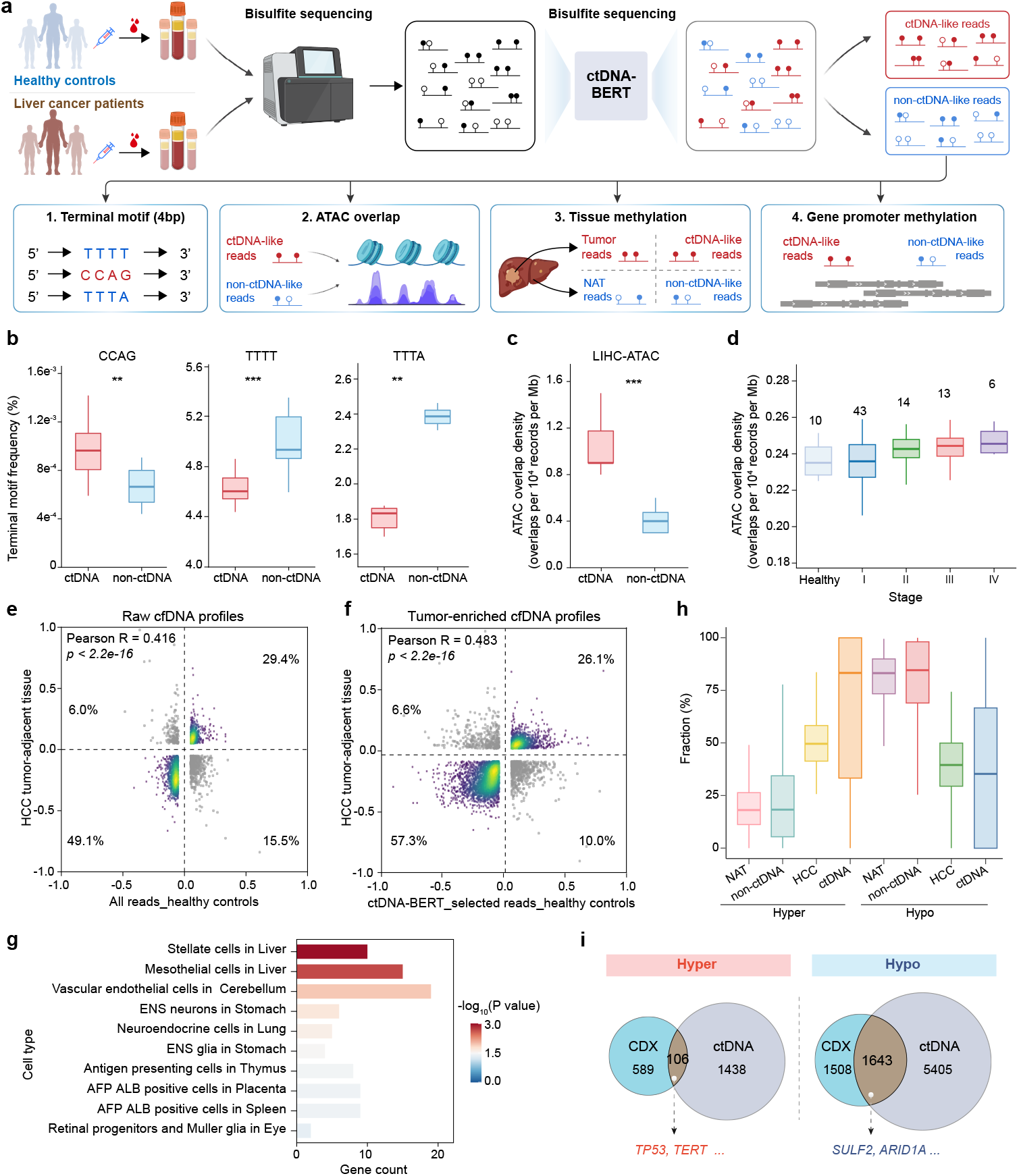
A frozen scorer enriches concordant molecular signals in independent HCC plasma. **a**, Workflow for 86 plasma samples (10 healthy controls and 76 HCC). The top 5% of reads formed the selected pool. **b**, Per-sample terminal 4-mer frequencies in selected and background pools. **c**, LIHC ATAC-seq overlap normalized per 10^4^ valid reads and per megabase of peak territory. **d**, Exploratory stage analysis using peaks selected for stage-associated behavior (healthy controls and stage I-IV HCC; n = 10, 43, 14, 13 and 6). Because the same samples defined and displayed the peak subset, this panel is exploratory and not directly comparable with c. **e,f**, CpG-island methylation differences between HCC tumor and adjacent tissue versus differences between HCC and control plasma, using all reads (e) or selected reads (f). Pearson r = 0.416 and 0.483, respectively (both P < 2.2 *×* 10^*−*16^). **g**, Cell-type categories over-represented among the top 250 genes; this analysis does not estimate cell fractions. **h**, Methylation distributions in independently defined hypermethylated and hypomethylated CpG sets. **i**, Promoter-level overlap between selected plasma reads and liver-cancer CDX. Box plots show medians, quartiles and 1.5 *×* interquartile-range whiskers. NAT, tumor-adjacent normal tissue; CDX, cell line-derived xenograft.

At CpG-island resolution, tissue–plasma methylation changes were positively correlated before and after read selection. Pearson *R* increased from 0.416 using all reads to 0.483 using selected reads (both P < 2.2 × 10^*−*16^; **Fig. 4e,f**), and directionally concordant islands rose from 78.5% to 83.4% while the discordant quadrant fell from 15.5% to 10.0%. CpG sets defined independently from HCC tumor versus adjacent tissue and from liver-cancer CDX versus WBC showed the same pattern (Fig. 4h and Supplementary Fig. 3a): selected reads followed tumor and CDX directionality, whereas background reads followed adjacent tissue and WBC directionality. At hypermethylated loci, the median methylated fraction was 83.3% in selected reads and 18.3% in background reads, compared with 49.6% in HCC tissue and 18.1% in adjacent tissue; hypomethylated loci showed the reciprocal pattern.

Among the 250 highest-ranked genes, the two strongest reference cell-type categories were liver stellate and mesothelial cells (**Fig. 4g**). Cell-type annotations for the expanded set of 500 gene candidates are shown in Supplementary Fig. 3b. A promoter-level comparison between selected plasma reads and liver-cancer CDX identified 106 shared hypermethylated and 1,643 shared hypomethylated genes (Fig. 4i), with overlap significance assessed against promoters with sufficient coverage in both datasets [56]. The hypomethylated set included *SULF2*, which is up-regulated in 57% of HCCs and associated with shorter survival [57], and *ARID1A*, a chromatin regulator recurrently mutated in liver cancer [58]. Promoter hypomethylation of *SULF2* is consistent with its reported up-regulation, whereas the functional consequence of *ARID1A* promoter hypomethylation was not assessed here.

### 2.5 A frozen scorer transfers to an independent stage III pan-cancer cohort without retraining

We tested transfer by applying the frozen scorer and the HCC annotation pipeline, without retraining or recalibration, to an independent multi-cancer plasma cohort (**Fig. 5**). The same chromatin-overlap, promoter-methylation and external-expression analyses were used unchanged.

**Fig. 5.**
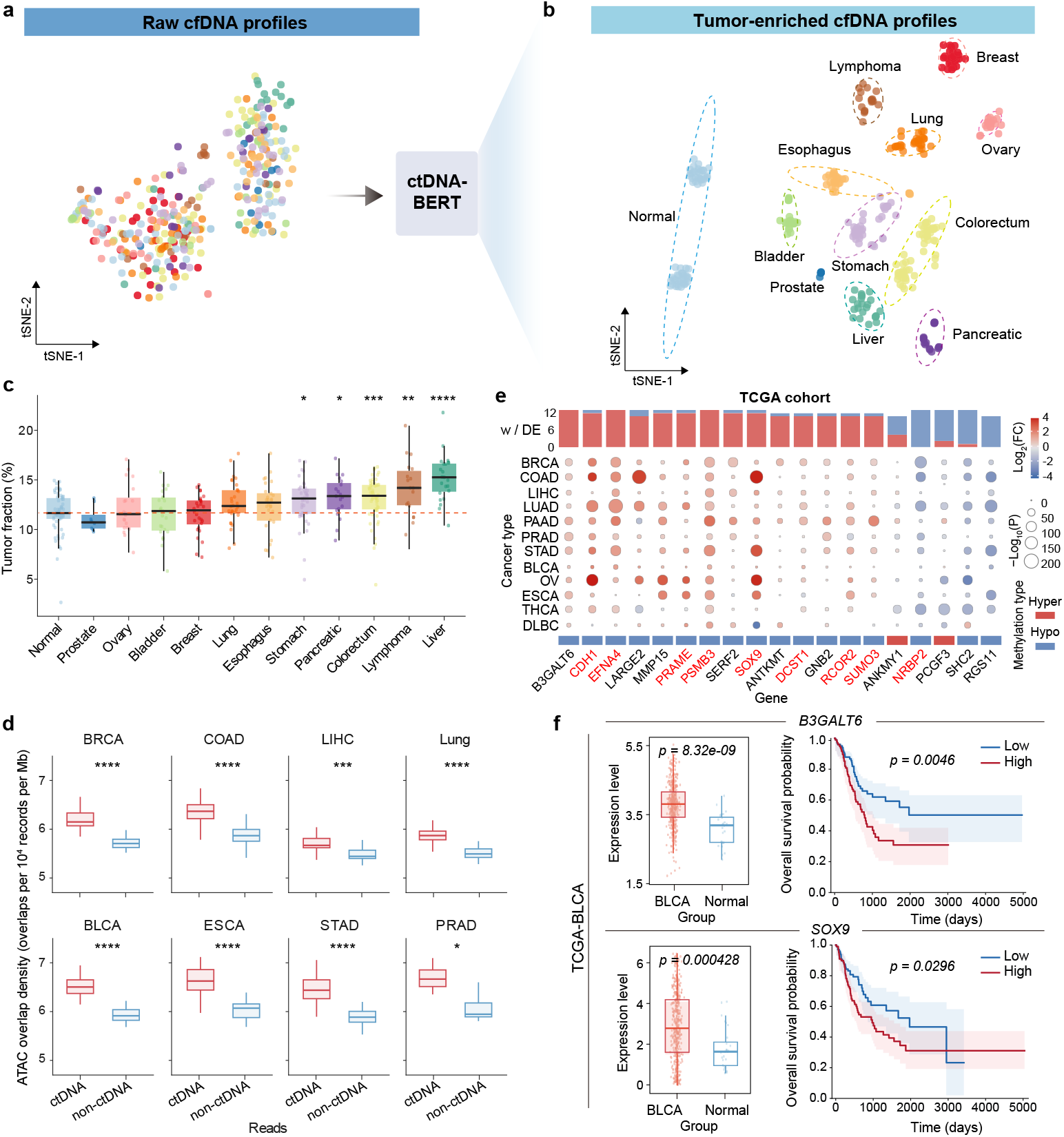
A frozen scorer transfers to an independent stage III pan-cancer cohort. **a,b**, t-SNE projections of CpG-island methylation profiles from 348 plasma samples, computed from all reads (a) and selected reads (b). The cohort comprised 58 healthy controls and 290 stage III cancers across 11 cancer types: breast (n = 28), colorectal (n = 54), liver (n = 23), lung (n = 30), non-Hodgkin lymphoma (n = 18), ovarian (n = 20), bladder (n = 23), esophageal (n = 28), gastric (n = 39), pancreatic (n = 19) and prostate cancer (n = 8). Adjusted Rand indices were computed in the full feature space. **c**, ctDNA-BERT-derived index by group. This analysis-specific quantity is not calibrated to an external clinical measurement. Asterisks indicate two-sided comparisons with healthy controls after Benjamini-Hochberg adjustment. **d**, ATAC-seq overlap of selected and background reads in eight cancer types, normalized per 10^4^ valid reads and per megabase of peak territory. **e**, Nineteen recurrent promoter candidates annotated in TCGA; dot size indicates −log10 P and color indicates log2 expression fold change. **f**, TCGA-BLCA expression and overall-survival associations for *B3GALT6* and *SOX9*. P values are unadjusted. THCA, thyroid; STAD, gastric; LUAD, lung adenocarcinoma; LIHC, liver; ESCA, esophageal; COAD, colorectal; BRCA, breast; BLCA, bladder.

The cohort comprised 290 stage III cancers across 11 cancer types and 58 healthy controls (348 specimens). Cancer cases included breast (n = 28), colorectal (n = 54), liver (n = 23), lung (n = 30), non-Hodgkin lymphoma (n = 18), ovarian (n = 20), bladder (n = 23), esophageal (n = 28), gastric (n = 39), pancreatic (n = 19) and prostate cancer (n = 8). Restricting this analysis to stage III disease separated the transfer evaluation from the stage I screening cohort.

Selected-read CpG-island profiles showed tumor-type grouping and separation of healthy controls not apparent in all-read profiles (**Fig. 5a,b**). In the full feature space, adjusted Rand index was 0.061 before selection and 0.223 after selection [59, 60]. The ctDNA-BERT-derived index was higher than in controls for 5 of 11 cancer types, with the largest differences in liver cancer and lymphoma (**Fig. 5c**).

Across eight cancer types, selected reads showed greater overlap with matched cancer-type ATAC-seq peaks than background reads ([53]; **Fig. 5d**). Overlap was normalized per 10^4^ valid reads and per megabase of peak territory. The peak resource, normalization and top-5% selection rule were carried over unchanged from the HCC analysis.

Promoter-level analysis identified 19 recurrent candidates: *B3GALT6, CDH1, EFNA4, LARGE2, MMP15, PRAME, PSMB3, SERF2, SOX9, ANTKMT, DCST1, GNB2, RCOR2, SUMO3, ANKMY1, NRBP2, PCGF3, SHC2* and *RGS11*. Seventeen were hypomethylated in selected reads; *ANKMY1* and *PCGF3* were hypermethylated (**Fig. 5e**). Additional promoter methylation comparisons are shown in Supplementary Fig. 4a,b. *PRAME*, a cancer-testis antigen activated by promoter hypomethylation, appeared in the expected direction [61]. In TCGA bladder cancer, *B3GALT6* expression differed between tumor and normal tissue (P = 8.32 × 10^*−*9^) and stratified overall survival (P = 0.0046). Corresponding P values for *SOX9* were 0.000428 and 0.0296 (Fig. 5f). *PCGF3* and *SUMO3* are shown in Supplementary Fig. 4c,d.

Thus, selected reads showed concordant chromatin and promoter annotations across independent cohorts, tumor types and analysis sites without cohort-specific adjustment.

### 2.6 Selected-read features separate stage I cancer from healthy controls and carry tissue-of-origin information across five cancer types

We next tested whether selected-read features distinguish stage I cancer from healthy controls. The retrospective case-control cohort comprised 730 stage I cancers (60 breast, 93 colorectal, 75 liver, 286 lung and 216 thyroid) and 747 healthy controls, totalling 1,477 specimens (**Fig. 6a**). No specimen contributed to pretraining or fine-tuning. Using the same chromatin annotation as in the HCC and pan-cancer cohorts, selected reads from all five cancers showed greater median overlap with matched ATAC-seq peaks than those from healthy controls (one-sided Mann-Whitney U tests with Benjamini-Hochberg adjustment; [53]; **Fig. 6b**), with separation strongest for thyroid, colorectal and lung cancers and weakest for liver cancer.

**Fig. 6.**
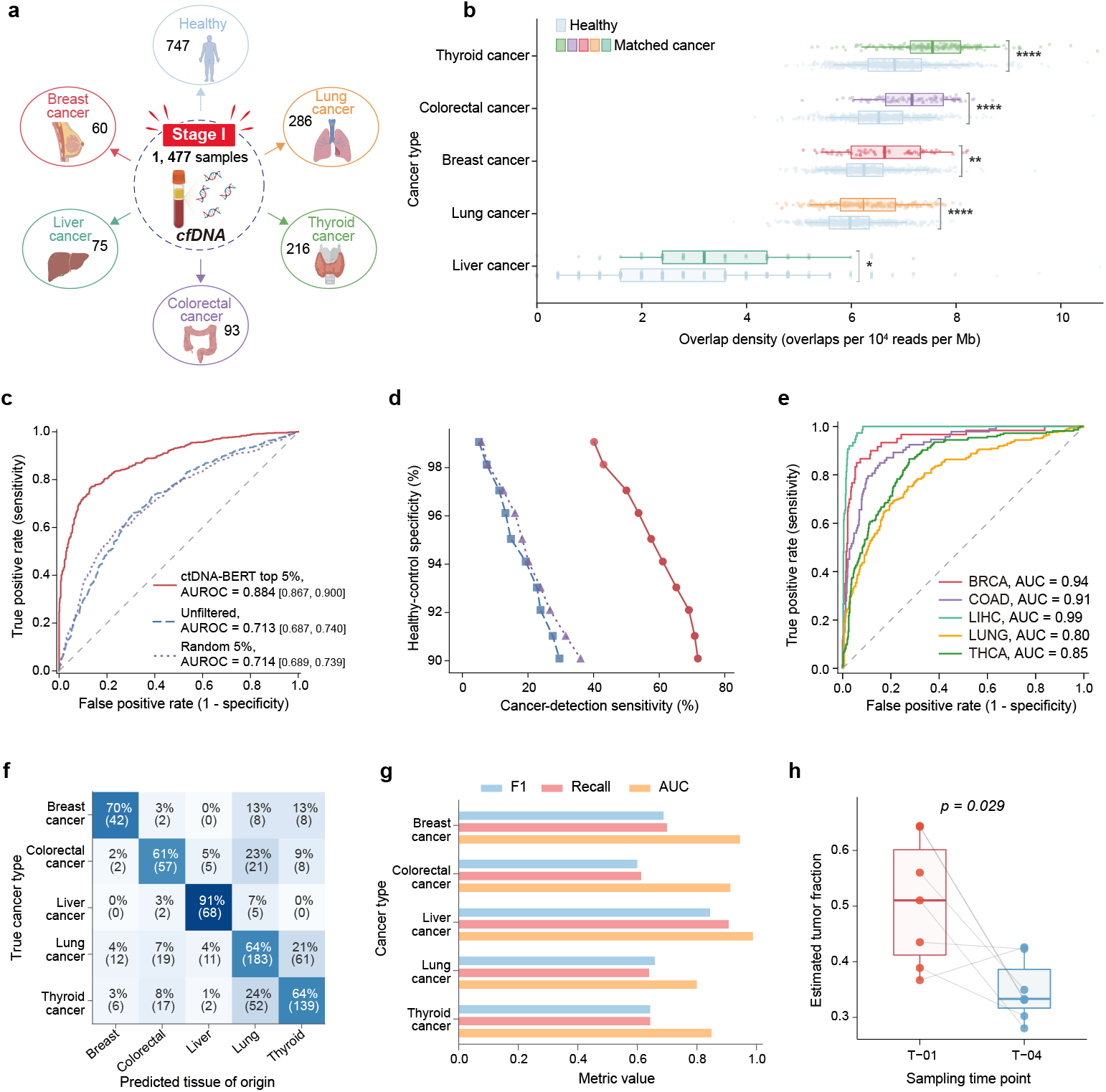
Selected-read features discriminate stage I cancer from control and carry tissue-of-origin information. **a**, Retrospective stage I case-control cohort (N = 1,477): 747 healthy controls and 730 cancers (60 breast, 93 colorectal, 75 liver, 286 lung and 216 thyroid). **b**, Cancer-matched ATAC-seq overlap in selected reads from each cancer type versus healthy controls, normalized per 10^4^ valid reads and per megabase of peak territory; one-sided Mann-Whitney U tests with Benjamini-Hochberg adjustment. **c**, Cancer-versus-control ROC curves for top-5%, unfiltered and random-5% read inputs, with displayed AUROC values of 0.884 [0.867, 0.900], 0.713 [0.687, 0.740] and 0.714 [0.689, 0.739], respectively. The selected-read model used fold-internal ANOVA selection of 2,000 of 5,489 inputs. **d**, Sensitivity-specificity trace from the same out-of-fold analysis as c, using the same feature-selection and SVM configuration. Thresholds were chosen post hoc from pooled scores. **e**, Nested out-of-fold one-versus-rest ROC curves. **f**, Five-class confusion matrix normalized by true cancer type. **g**, Class-specific F1, recall and AUC. **h**, Paired comparison of the ctDNA-BERT-derived index between T-01 and T-04 in seven patients with colorectal cancer (two-sided paired test, P = 0.029). Box plots show medians, quartiles and 1.5 *×* interquartile-range whiskers.

The selected-read classifier was evaluated by five-fold stratified cross-validation, with each specimen scored outside its training fold. Figure 6c also compares the ctDNA-BERT-selected input with unfiltered-read and randomly selected 5%-read inputs. The model combined 5,456 k-mer frequencies with 33 BED-derived annotations; within each training fold, ANOVA selection retained 2,000 variables, which were then scaled and used to fit a class-balanced radial-basis-function support-vector machine. Feature selection, scaling and model fitting were confined to the training folds. The displayed AUROC was 0.884 [0.867, 0.900] for the ctDNA-BERT top-5% input, compared with 0.713 [0.687, 0.740] for unfiltered reads and 0.714 [0.689, 0.739] for random 5% selection (Fig. 6c). Figure 6d summarizes the same out-of-fold analysis, with sensitivity ranging from 40.1% at 99.1% specificity to 71.8% at 90.1% specificity. In a separate nested five-fold analysis of the 730 cancers, one-versus-rest AUCs were 0.94 for breast, 0.91 for colorectal, 0.99 for liver, 0.80 for lung and 0.85 for thyroid cancer (**Fig. 6e**); recalls were 70%, 61%, 91%, 64% and 64%, respectively (**Fig. 6f,g**). The largest reciprocal confusion occurred between lung and thyroid cancer, with 61 of 286 lung cases assigned to thyroid and 52 of 216 thyroid cases assigned to lung.

Seven patients with colorectal cancer contributed serial specimens. In six of the seven, the ctDNA-BERT-derived index was lower at T-04 than at T-01 (two-sided paired test, P = 0.029; **Fig. 6h**). T-02 and gene-level promoter methylation summaries are shown in Supplementary Fig. 5; individual-patient promoter coverage and methylation comparisons are shown in Supplementary Fig. 6.

## 3 Discussion

What this work contributes is a layer rather than a classifier. cfDNA analysis has conventionally compressed molecules into regional or sample-level summaries before any inference is attempted; ctDNA-BERT inserts a scoring step ahead of that compression, assigns every eligible molecule a tumor-like score from a single representation that jointly encodes nucleotide context, CpG state and a strand-normalized fragment end, and then exposes that score for arbitrary downstream use. Two obstacles had to be removed before such a layer was possible, and each is methodological rather than incremental: where molecule-level supervision can come from, and how molecular information can survive into the representation at all. Matched ablations show that masked-language pretraining, the paired terminal segment and methylation-aware encoding each contribute to tumor-background separation. Attention analyses show layer-dependent weighting of methylation-bearing and sequence tokens. The main comparison includes *E2F2, E2F4, FOXC1, ELF3, YY1, TEAD1, TCF7L2, PPARG, POU5F1* and *FOS* relative to *CTCF* (Fig. 2f), with complementary *LEF1* and *E2F1* examples in Supplementary Fig. 1c. The *TERT* /*E2F2* display and supplementary locus examples provide genomic context for attention weights, rather than evidence of TF occupancy or causal regulation (Fig. 2g; Supplementary Fig. 1d,e)[53]. The representation was validated by three convergent benchmarks: synthetic bulk mixtures, held-out xenograft mixtures carrying molecule-level ground truth, and an orthogonal pathologist-assigned FFPE tumor-content series. It was then tested by transfer to an independent stage III pan-cancer cohort and to a stage I-restricted case-control cohort in which tumor signal is scarcest.

The framework differs from atlas-style methods that aggregate methylation before inference and from DMR-gated read-level models such as DISMIR and MethylBERT [31, 32]. Recent read-level studies have established the value of scoring individual molecules, so the contribution here is not read-level modeling itself. It is, first, that the supervision is obtained by construction rather than by inference: xenograft dual-genome alignment yields molecules whose tumor origin is fixed by the experimental design, which removes the circularity of training a molecular model on labels derived from the regional statistics that model is meant to replace. It is, second, that methylation enters the token alphabet and the fragment end enters as a paired segment, so that epigenetic state, nucleotide context and terminal position interact inside a single molecular representation instead of being reconciled after each has already been averaged away. DMR independence and cross-cohort transfer are consequences of these two choices rather than separate claims standing alongside them. It also addresses a different problem from Transformer-based approaches that couple raw-read representations directly to sample-level multiple-instance prediction [35–38]. Those methods optimize an end-to-end sample-level classifier; ctDNA-BERT instead exposes a fixed, frozen molecular score that can be inspected, thresholded and reused independently of any particular downstream sample model. This separation of molecular scoring from sample-level inference, rather than the use of a Transformer architecture alone, is the central methodological distinction.

This separation also broadens how the scorer can be used. Without retraining or recalibration, the same frozen checkpoint transferred across three independent applications: HCC plasma, an independent stage III pan-cancer cohort and a retrospective stage I case-control cohort. Each converged on molecular annotations that were never available during training, including fragment-end composition, chromatin accessibility, tissue-level methylation direction and promoter-level tumor concordance. A scoring layer that transfers unchanged across cohorts and cancer types could precede existing cfDNA methylation workflows without replacing them, supporting mean scores, thresholded rates or ranked read pools as different analyses require. Enriched pools may reduce the read depth needed for downstream molecular annotation and could serve as a front-end selection step in minimal-residual-disease-style workflows [45], but these applications require dedicated prospective testing.

Several limitations define the scope of the current evidence. Clinical interpretation is limited by the retrospective case-control design, the small paired longitudinal series (n = 7) and the absence of benign or inflammatory disease controls alongside healthy controls. The xenograft label is also specific in what it certifies. It establishes that a read is tumor-derived, and it does so without circularity, but xenograft tumors are propagated in a murine host and the fragments they release need not reproduce every property of circulating tumor DNA, in particular the nuclease processing that shapes fragment ends in human plasma. The frozen scorer is therefore supervised by molecules whose tumor identity is certain and whose fragmentation context is only approximate; the plasma analyses in Figs. 4 to 6 are the evidence that the resulting score nonetheless transfers, and a direct comparison of xenograft-derived and plasma-derived terminal-motif distributions would further bound this assumption. Future work should prospectively lock preprocessing, scoring and aggregation procedures ahead of evaluation, add benign and inflammatory controls, assess calibration across sequencing and library-preparation protocols, and validate performance across independent centers and populations [48, 62].

## 4 Online Methods

### 4.1 Study design

We evaluated ctDNA-BERT as a read-level computational framework for bisulfite-sequencing data. The three training stages and all downstream benchmarks were separated at the specimen level. The independent HCC cohort, the stage III pan-cancer cohort and the stage I case-control cohort contributed to none of the three training stages.

Training proceeded in three steps, referred to throughout by name rather than by number so that they are never confused with clinical stage. Pretraining is self-supervised masked-language modeling. Tissue fine-tuning is supervised training that separates tissue-derived reads from the hematopoietic-background class, denoted HEP. Xenograft fine-tuning is supervised training that separates human xenograft tumor reads from white-blood-cell reads. Parameters were frozen after xenograft fine-tuning for the specified plasma applications. The organ-system tasks and FFPE-specific fine-tuning are distinct model branches, and sample-level classifiers were fitted within their designated training folds. We reserve the term calibration for procedures that map model outputs onto an externally measured quantity; in this study that applies only to the FFPE tumor-content analysis, where predictions were mapped onto pathologist-assigned tumor content. The three training stages update parameters and are therefore described as pretraining and fine-tuning throughout.

The organ-system evaluation in Fig. 3e comprises seven independent binary tasks initialized from the pretraining checkpoint at 400,000 optimization steps, and is interpreted separately from applications of the frozen scorer.

### 4.2 Data inventory and partitioning

The dataset inventory comprises 415 Primer-RRBS libraries for pretraining; a 463-library study tissue inventory and 177 public RRBS reference libraries for tissue fine-tuning; specimen-separated xenograft material and derived benchmark mixtures for xenograft fine-tuning; and independent FFPE, HCC, stage III pan-cancer, stage I case–control and serial cohorts. Table 1 maps each dataset family to its analytical role, verified inventory and access category.

**Table 1.** Dataset inventory, analytical role and access route.

| Dataset family | Verified inventory | Analytical role | Source and access category |
| --- | --- | --- | --- |
| **Pretraining libraries** | 415 -Primer-RRBS libraries | masked-language pretraining | Study-generated human plasma cfDNA |
| **Tissue fine-tuning inventory** | 463 libraries: 196 normal; 267 tissue- or cancer-labeled | tissue fine-tuning | Study-generated human tissue |
| **Public HEP reference panel** | 177 libraries from 14 GEO series | Hematopoietic-background class; Fig. 3e comparator | Public GEO data |
| Higher-fraction six-source WGBS/RRBS mixtures | 6 tumor source categories; 9 target fractions spanning 10–90%; 3 composition regimes (Uniform, Dirichlet and Constrained) per platform | 10–90% benchmark; Supplementary Fig. 2a–d | Study-generated RRBS in silico mixtures |
| Low-fraction WGBS/RRBS mixtures | 19 distinct target fractions spanning 0.1–10%; 30 mixtures per target per platform; 570 derived mixtures per platform | Fig. 3a lower panels; platform-specific inputs and metrics | Study-generated RRBS in silico mixtures |

**Table 1** Dataset inventory, analytical role and access route. (continued)
| Dataset family | Verified inventory | Analytical role | Source and access category |
| --- | --- | --- | --- |
| **Xenograft mixtures and rare-read series** | 3 PDX cancer types (COAD, LUAD, STAD); 285 low-fraction mixtures (19 targets, 0.1–10%; 5 replicates); 270 higher-fraction mixtures (9 targets, 10–90%; 5 replicates); 5 rare-read levels ($10^{-1}$–$10^{-5}$) | Xenograft fine-tuning; held-out benchmarks (Fig. 3b–d; Supplementary Fig. 2e) | Study-generated PDX/CDX data |
| **Organ-system binary tasks** | 65 cancer specimens from 12 types + 177 HEP libraries | Seven system-versus-HEP tasks (Fig. 3e) | Study-generated cancer data and public HEP data |
| **FFPE tumor-content cohort** | 31 specimens: 23 non-liver development; 8 independent liver test | Fine-tuning and calibration on 23; frozen test on 8 (Fig. 3f–h) | Study-generated human tissue |
| **Independent HCC plasma** | 86 samples: 10 healthy; 76 HCC | Frozen-scorer molecular concordance (Fig. 4) | Study/collaborator human data |
| **Stage III pan-cancer plasma** | 348 samples: 58 healthy controls; 290 stage III cancers across 11 cancer types | Frozen-scorer transfer and annotation (Fig. 5) | Study/collaborator human data |
| **Stage I case-control and serial plasma** | 1,477 specimens: 730 stage I cancers (60 breast, 93 colorectal, 75 liver, 286 lung, 216 thyroid); 747 healthy controls; 7 serial CRC cases | Binary, tissue-of-origin and longitudinal analyses (Fig. 6) | Study/collaborator human data |
| **Public post-inference annotations** | ATAC-seq peaks, motif intervals, TCGA matrices, methylation atlas and hg19 tracks | Biological annotation and downstream classifier features | Cited public resources |

The HEP reference panel contains 177 RRBS libraries from 14 NCBI Gene Expression Omnibus series: GSE105127, GSE114254, GSE114807, GSE118255, GSE130582, GSE135770, GSE151527, GSE160331, GSE164365, GSE165360, GSE59163, GSE66167, GSE71702 and GSE93383. The study tissue inventory comprised 463 libraries, including 196 labeled normal and 267 tissue- or cancer-labeled libraries. Five libraries, one from each of the breast, liver, lung, pancreas and thyroid groups, were reserved for testing at the tissue fine-tuning stage. The remaining 458 libraries were used for tissue fine-tuning, and their organ-site composition is shown in Fig. 1c (middle panel).

### 4.3 Bisulfite-sequencing preprocessing

WGBS and Primer-RRBS reads were aligned to the hg19 reference genome and methylation calls were generated with the project-specific Bismark workflow [51]. Unmapped and low-quality reads, PCR duplicates, reads overlapping the project blacklist and reads failing assay-specific quality filters were removed. Sample identifiers were preserved throughout preprocessing so that dataset partitioning was completed before read sampling. Molecular sequences were limited to 200 bases before subword tokenization. After tokenization each input was truncated or padded to a maximum sequence length of 64 tokens, with padded positions excluded by the attention mask.

As in conventional reduced-representation bisulfite sequencing [46], the Primer-RRBS protocol constrains one fragment end while retaining greater sequence variability at the opposite end. The preprocessing workflow extracted the variable 3^*′*^ terminal 6-base sequence according to the aligned strand. Molecular sequences and terminal sequences from minus-strand reads were transformed to a common orientation before tokenization. The terminal 6-base sequence was then encoded as a paired input segment with its own segment identity rather than being appended as a scalar feature after sequence encoding.

### 4.4 Human-read isolation from xenografts

Patient-derived xenograft (PDX) and cell line-derived xenograft (CDX) libraries contain both human tumor-derived and murine host-derived reads. Reads were evaluated against the hg19 and mm9 reference genomes, and only those uniquely assigned to hg19 and passing sequence-quality filters were retained as human tumor-derived material. Xenograft specimens, rather than individual reads, were assigned either to xenograft fine-tuning or to held-out mixture construction, preventing specimen overlap between the two datasets. White-blood-cell reads served as the common background component in both settings.

### 4.5 Methylation-aware token strings and paired terminal input

For read *i*, let 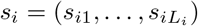 denote the observed sequence, 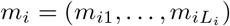 the aligned CpG-state vector, *e*_*i*_ the terminal 6-base sequence and *η*_*i*_ ∈ {+, −} the aligned strand. The preprocessing map places the molecular and terminal sequences in a common orientation:

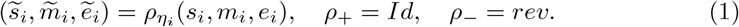

Here, *rev* denotes the strand-dependent transformation applied before tokenization. Let *c*_*ij*_ indicate whether oriented position *j* begins a CpG context.

Methylation-aware encoding was defined as

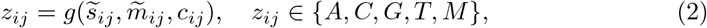

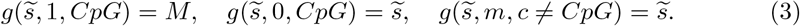

Methylated CpG cytosines therefore enter the discrete molecular alphabet as M, whereas unmethylated CpGs and non-CpG positions retain their oriented sequence symbols. Because M is a symbol in the alphabet rather than an auxiliary feature, methylation state is available to the tokenizer and to every subsequent encoder operation.

After orientation normalization, M-containing strings yield MG or GM units during subword tokenization; GM arises where a subword merge places the guanine of a CpG before the methylated cytosine of the following dinucleotide context. We trained a SentencePiece tokenizer using the byte-pair encoding algorithm on these molecular strings [63, 64]. The base vocabulary contained 4,096 entries; adding MG and GM produced the 4,098-entry vocabulary used for training. Learned subword units rather than fixed-length k-mers follow current practice for DNA language models [49, 50]. The tokenizer configuration supplied the special tokens.

Let *T*(·) denote subword tokenization and ⊕ sequence concatenation. The paired input was

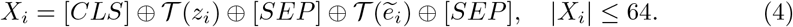

The segment vector 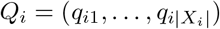 assigns *q*_*ij*_ = 0 to fragment-body tokens and *q*_*ij*_ = 1 to terminal tokens. Terminal sequence therefore participates in the same self-attention operations as nucleotide and methylation-containing tokens, while segment identity remains available to the ctDNA-BERT encoder.

### 4.6 ctDNA-BERT encoder and read-level probability

ctDNA-BERT used a bidirectional BERT-style encoder to model sequence context on both sides of each CpG. [65, 66]. The principal configuration had *L* = 12 layers, hidden size *d* = 768, *H* = 12 attention heads, head dimension *d*_*h*_ = *d/H* = 64, feedforward size 3,072, hidden and attention dropout of 0.1, a maximum sequence length of 64 tokens and two segment types. For token matrix *X*_*i*_ and segment vector *Q*_*i*_, the initial hidden states were

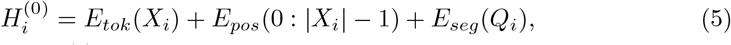

where each row of 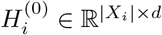 represents one input position. Head *h* in layer *l* projects queries, keys and values from the preceding hidden states:

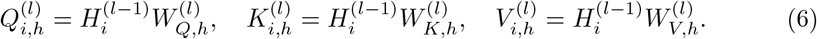

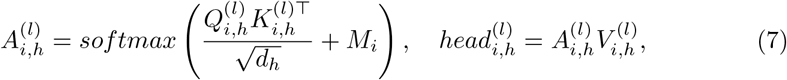

where *M*_*i*_ masks padded key positions with a large negative value. The heads were concatenated and passed through the output projection, residual path and layer normalization:

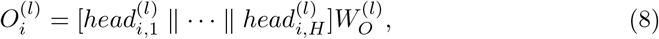

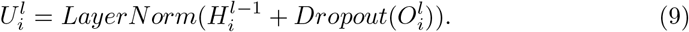

The position-wise feed-forward transformation and second residual path were

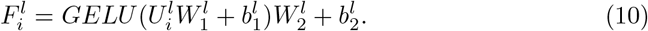

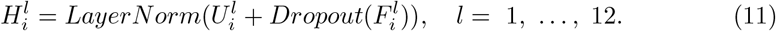

Because MG and GM have distinct token embeddings, methylation state changes the projected queries, keys and values at every layer. For fixed tokenization, the query-side change in the first-layer attention logit between position j and context position k is

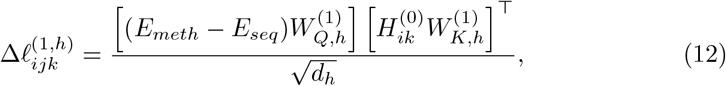

where *E*_*meth*_ and *E*_*seq*_ denote the position-*j* token embeddings under methylation-aware and sequence-only encoding, respectively. An analogous key-side term applies when attention is directed toward the methylation-containing position. Methylation state therefore perturbs contextual encoding before the classification head.

The final [CLS] state was passed to a classification head with two output units. Because trained configurations use different numeric label indices, let o(i,T) and o(i,B) denote the tumor and background logits by name rather than by index. The read-level tumor probability was

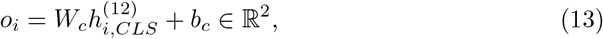

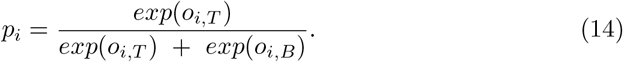

During xenograft fine-tuning the named tumor class denotes human xenograft material and the background class denotes white-blood-cell material. In patient plasma the resulting softmax value is termed a tumor-like score; it does not establish the cellular origin of an individual read and does not guarantee cross-assay comparability.

### 4.7 Training curriculum

Pretraining used masked language modeling, with 15% of token positions selected for prediction. For masked-position set M_*i*_ and corrupted input 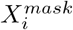, the reconstruction loss was

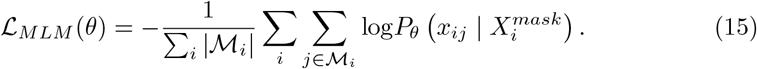

Pretraining used AdamW for 10 epochs with a learning rate of 1 × 10^*−*5^, per-device train and evaluation batch sizes of 256, weight decay 0.01, linear learning-rate scheduling, bfloat16 mixed-precision arithmetic, no warm-up and seed 42 [67, 68]. The checkpoint at 460,000 optimization steps initialized tissue fine-tuning. In accordance with the Fig. 1 schematic, tissue fine-tuning used the binary class set C = {tissue, HEP}. Its cross-entropy objective was

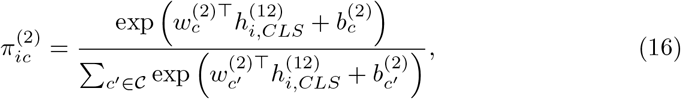

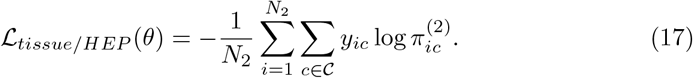

Xenograft fine-tuning transferred the ctDNA-BERT encoder to a xenograft-versus-white-blood-cell objective. Using tumor probability *p*_*i*_, the binary objective was

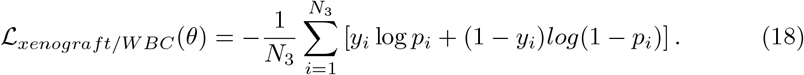

The parameter-transfer path was

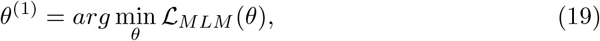

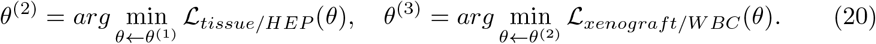

Xenograft fine-tuning used AdamW, a learning rate of 2 × 10^*−*5^, per-device training and evaluation batch sizes of 512, weight decay 0.01, linear scheduling, warm-up ratio 0.01, bfloat16 mixed-precision arithmetic and seed 42. Evaluation and check-pointing occurred every 10,000 steps, and the checkpoint with the lowest evaluation loss was frozen for all downstream scoring. Implementation used PyTorch and the Transformers library [69, 70].

### 4.8 Probability aggregation and read selection

For sample *s*, let ℛ_*s*_ denote the eligible reads and *n*_*s*_ = |ℛ_*s*_|. We summarized read probabilities by their mean and by the proportion above threshold *τ* :

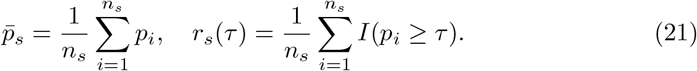

For analyses requiring equal read depth, score rank defined the retained set:

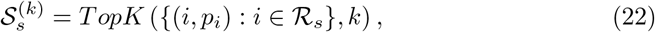

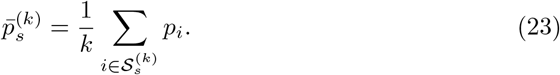

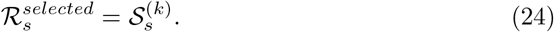

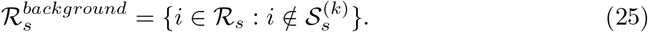

The neural scoring function was frozen after xenograft fine-tuning. Downstream summaries used the prespecified mean, threshold or rank rule appropriate to each analysis. Table 2 records the read partition and sample-level aggregation used by each main-figure analysis. Quantities labeled as a ctDNA-BERT-derived index remain specific to their corresponding estimator and are not externally calibrated tumor fractions.

**Table 2.** Panel-level read selection and aggregation rules.

| Panel | Read filtering | Threshold, k or tie rule | Minimum eligible reads | Sample-level aggregation |
| --- | --- | --- | --- | --- |
| **Fig. 1c** | Preprocessing-qualified reads | None | One read | Read-level softmax tumor probability |
| **Fig. 2a** | Common held-out labeled evaluation reads | None; threshold-free | Common evaluation set at each step | Read-level ROC AUC per configuration and step |
| **Fig. 2b** | Same evaluation reads | None | Same set | k-means ($k = 2$) on final-layer [CLS] representations in 768-d space; ARI against read labels |
| **Fig. 2c,d** | Evaluation reads with eligible token pairs | None | At least one pair in the summarized category | Mean across heads and pairs within read, then median across reads |
| **Fig. 2f,g** | Eligible reads intersecting the stated motif interval | No score-rank selection | At least one motif-associated token | Maximum token attention within motif; single-read display for g |
| **Fig. 3a-c** | All eligible reads in each constructed mixture | No selected/background partition | 30 million requested reads per mixture | Analysis-specific fraction estimate from read probabilities |
| **Fig. 3d** | Labeled tumor and background reads at the stated dilution | None | Defined by each dilution set | Read-level ROC AUC |
| **Fig. 3e** | Reads entering each independently trained binary head | Larger of two class logits | No additional minimum | Specimen-level binary accuracy |
| **Fig. 3f-h** | Eligible reads from pathologist-delineated FFPE regions | Frozen FFPE calibration mapping | One evaluable specimen | Calibrated specimen tumor-content estimate; R2 and Spearman rho |
| **Fig. 4b-i** | Eligible HCC-cohort reads | Top 5% within sample by tumor probability; stable order, exactly k retained | At least one retained read for the reported modality | Per-sample motif frequency, methylation fraction or normalized overlap |

**Table 2** Panel-level read selection and aggregation rules. (continued)
| Panel | Read filtering | Threshold, k or tie rule | Minimum eligible reads | Sample-level aggregation |
| --- | --- | --- | --- | --- |
| **Fig. 5a,b,d-f** | Eligible stage III cohort reads | Top 5% within sample by tumor probability | One evaluable sample | Per-sample selected-read or normalized annotation summaries; ARI for a,b |
| **Fig. 5c** | Eligible stage III cohort reads | Analysis-specific aggregation | One evaluable sample | ctDNA-BERT-derived index (not an externally calibrated tumor fraction) |
| **Fig. 6b** | Eligible stage I cohort reads | Top 5% within sample | One evaluable sample | Per-sample normalized ATAC overlap |
| **Fig. 6c-g** | Eligible stage I cohort reads | Top 5% within sample; classifier selection is fold-internal | One complete sample feature vector | Out-of-fold SVM score or nested out-of-fold class probability |
| **Fig. 6h** | Eligible serial CRC reads | Frozen serial-analysis estimator | One evaluable specimen per time point | ctDNA-BERT-derived index; paired T-01 versus T-04 comparison |

### 4.9 Ablation and representation analyses

We compared the complete ctDNA-BERT configuration with three matched variants, each omitting exactly one element: the pretraining checkpoint, the paired terminal segment, or methylation-aware encoding. All four variants used identical training schedules, identical data and the same held-out set of labeled evaluation reads, and were scored at 0, 4,000, 8,000, 12,000, 16,000 and 20,000 optimization steps. Because every evaluation read carries a tumor or background label, class separation was summarized by read-level ROC AUC at each step rather than by the shape of the score distributions.

Final read representations were compared by taking the last-layer [CLS] representation of each evaluation read, clustering those representations with k-means (k = 2, seed 42) in the full 768-dimensional embedding space, and scoring the clustering against the true read labels with the adjusted Rand index [60, 71]. Clustering was performed in the embedding space and not on two-dimensional coordinates, because t-SNE preserves local neighborhoods but not global distances, so a partition induced on its output reflects the projection as well as the representation [59, 72]. t-SNE projections are shown for visual inspection only and were computed with the same procedure and perplexity for all variants [73].

Attention weights were exported from every encoder layer. Token pairs were classified as methylation-bearing to methylation-bearing, mixed or sequence-only. Let *P*_*i,c*_ denote token pairs in read *i* assigned to category *c*. Attention was first averaged across heads and eligible pairs within each read,

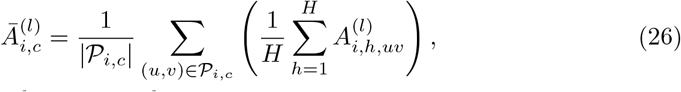

and then summarized across reads:

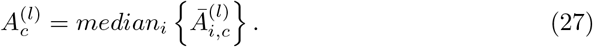

This two-level aggregation prevents reads with more eligible pairs from dominating the layer estimate. Because the three pair categories are measured on the same reads, layer-12 comparisons between categories used paired two-sided Wilcoxon signed-rank tests with Benjamini–Hochberg adjustment rather than unpaired tests.

For motif analysis, eligible LIHC reads were intersected with UCSC annotations and the JASPAR 2024 CORE vertebrate collection [52], and the maximum token attention within each motif interval was summarized. *CTCF* was specified in advance as the reference because it is the most frequently annotated motif among the eligible intervals. Motifs were compared with *CTCF* using two-sided Mann–Whitney U tests followed by Benjamini–Hochberg adjustment. Attention at methylation-bearing tokens and at motif-associated sequence tokens was evaluated separately; no adjacency, continuity or overlap criterion between CpG and motif positions was imposed. Single-read displays retained genomic context, motif identity, promoter annotation and token-level attention. Attention weights describe how the model distributes weight over its own input; they are not a faithful attribution method and were not interpreted as occupancy or causality [74–76].

### 4.10 Multi-source and xenograft mixture benchmarks

The six-cancer bulk-RRBS benchmark combined human tumor-tissue reads from liver, lung, colon, pancreas, ovary and prostate with white-blood-cell and other background reads. For the higher-fraction RRBS benchmark, *τ* ∈ {0.1, 0.2, …, 0.9} denotes the total tumor fraction on the 0–1 scale, *w* = (*w*_1_, …, *w*_6_) the normalized tumor-source weights, and *v* = (*v*_1_, …, *v*_*B*_) the normalized background weights. For the lower panels of Fig. 3a,b, the distinct target fractions on that scale are {0.001, 0.002, 0.003, 0.004, 0.005, 0.006, 0.007, 0.008, 0.009, 0.010, 0.020, 0.030, 0.040, 0.050, 0.060, 0.070, 0.080, 0.090, 0.100}; the shared 1% boundary is counted once in the combined grid. The target fraction assigned to tumor source k and background component b was

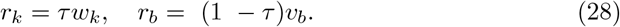

Three composition regimes were evaluated, because spatial structure and subclonal architecture make the tumor contribution to cfDNA unequal across regions and lesions [77]. Uniform mixtures assigned w_*k*_ = 1/6. Dirichlet mixtures used normalized weights drawn from a symmetric Dirichlet distribution with *α*_*k*_ = 0.05 for each of the six tumor sources, producing sparse and strongly unequal compositions; the normalized Dirichlet weight matrix was used without modification. Constrained mixtures designated one tumor source as dominant, with a nominal target of approximately 80% of the tumor component and the remaining sources contributing smaller fractions; the normalized constrained vectors were used without modification and their realized dominant weights ranged from 0.621 to 0.740. The three regimes therefore modeled balanced shedding, heterogeneous or long-tailed shedding across regions or lesions, and a dominant primary lesion accompanied by weaker secondary signals.

Each mixture targeted *N*_0_ = 30,000,000 reads. For component BAM *c* with *M*_*c*_ mapped reads and target fraction *r*_*c*_, the requested read count and BAM-level subsampling fraction were

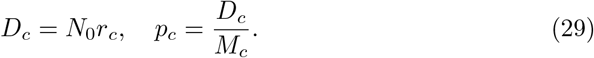

For the low-fraction WGBS and RRBS benchmarks, mixtures were generated separately at 19 distinct target tumor fractions: 0.1–1.0% in increments of 0.1 percentage points and 1–10% in increments of 1 percentage point, with the shared 1% target counted once. Each platform included six source-composition configurations (D1– D6), with five in silico downsampling replicates per configuration and target fraction, yielding 30 mixtures per target and 570 derived mixtures per platform. Mapped component counts were obtained with samtools idxstats, and the requested subsampling fraction *r*_*c*_*N*_0_*/M*_*c*_ was rounded to six decimal places [78]. Component BAMs were subsampled, indexed, merged, coordinate-sorted and re-indexed. WGBS comparisons included ctDNA-BERT, DISMIR, methAtlas and UXM, whereas RRBS comparisons included ctDNA-BERT, UXM, methAtlas, cfSort and CelFEER. Analyses used the available outputs for each method; the constructed mixture counts do not imply identical analyzed counts across methods [2, 3, 14, 15].

The xenograft benchmark combined held-out human tumor reads from colorectal, lung and gastric PDX specimens with a common white-blood-cell background. For target tumor-derived fraction *π*, the requested component fractions were

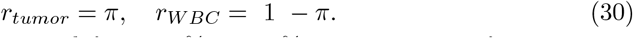

Target tumor fractions ranged from 0.1% to 1.0% in increments of 0.1 percentage points and from 1% to 10% in increments of 1 percentage point, yielding 19 distinct target fractions. The xenograft mixtures used the same 30-million-read target, component-read counting, six-decimal subsampling calculation, five-seed schedule, merge, coordinate sort and indexing procedure as the multisource mixtures. ctDNA-BERT, MethylBERT and DISMIR analyzed identical mixture files [31, 32]. Recovery was summarized using R^2^, root-mean-square error, mean absolute error and absolute-error distributions, reported per tumor type and per composition regime rather than pooled, because the ranking of methods differs between regimes. Separately, read-level ranking was assessed after downsampling labeled tumor reads to proportions of 10^*−*1^, 10^*−*2^, 10^*−*3^, 10^*−*4^ and 10^*−*5^. ROC AUC is invariant to class prevalence, so these values characterize ranking behavior and do not define a sample-level limit of detection; a precision–recall summary is the appropriate complement at extreme imbalance [54, 79].

Figure 3e summarized seven independent organ-system-versus-HEP binary evaluations initialized from the pretraining checkpoint at 400,000 optimization steps; it is not a seven-class organ-system classifier. The analysis included 65 cancer specimens from 12 cancer types, pooled as lymphatic (lymphoma, n = 5), respiratory (lung, n = 10), digestive (esophageal, colorectal, gastric and liver, n = 10 specimens in total), 30 reproductive (ovarian, n = 10), urinary (kidney, n = 10), endocrine (thyroid and pancreatic, n = 10 in total) and exocrine (prostate and breast, n = 10 in total). Each pooled group was compared separately with the same 177-library HEP reference panel. Accuracies were 0.930, 0.909, 0.889, 0.879, 0.869, 0.829 and 0.791 respectively. The lymphatic estimate rests on five specimens and is correspondingly imprecise.

### 4.11 FFPE tumor-content evaluation

The FFPE collection comprised colorectal cancer (n = 8), gastric cancer (n = 8), lung cancer (n = 7) and liver cancer (n = 8) specimens. A pathologist delineated tumor and adjacent non-tumor regions on H&E-stained FFPE sections and assigned tumor content to the annotated region. Corresponding adjacent-section material supplied the sequencing input (Fig. 3f). Reads from the tumor and adjacent non-tumor regions of the 23 colorectal, gastric and lung specimens were used to fine-tune ctDNA-BERT from the frozen scorer for binary tumor-versus-adjacent-tissue classification. The pathologist-assigned tumor-content values for those 23 specimens were then used to calibrate the model predictions and derive FFPE tumor-content estimates. This is the one calibration step in the study, in the sense defined above: model outputs were mapped onto an externally measured quantity [55].

After fine-tuning and calibration the ctDNA-BERT parameters and the calibration mapping were frozen and applied without further adjustment to an independent liver-cancer FFPE cohort of eight specimens, none of which had contributed to fine-tuning or calibration. MethylBERT and DISMIR used the same stated 23-specimen non-liver development cohort and eight-specimen liver test cohort. Their fine-tuning and calibration were performed separately for each method, and their combined results are shown in Fig. 3h. Agreement with the pathology reference was summarized by the R^2^ values shown in Fig. 3g,h. Each test observation was an independent FFPE specimen (n = 8); these R^2^ values do not provide a separate confidence interval for the agreement statistic. The pathology reference is itself a visual estimate with documented inter-observer variability, which bounds the agreement any method can demonstrate against it [80, 81]. This analysis evaluated tumor-content estimation in fixed tissue and was not used to calibrate plasma ctDNA fractions.

### 4.12 HCC plasma analyses

We applied the frozen scorer to 86 plasma samples comprising 10 healthy controls and 76 HCC cases. Within each sample the top 5% of reads by tumor probability formed the selected pool and the remainder formed the background pool. Terminal 4-base frequencies were calculated separately for the two pools and normalized by the number of valid reads in each. Overlap with LIHC ATAC-seq peaks was normalized per 10^4^ valid reads and per megabase of peak territory [53]. Terminal-motif frequencies, CpG-set fractions and ATAC overlap were independent summaries of the two pools; no analysis required two features to occupy the same read or genomic position.

The exploratory stage panel used an LIHC peak subset selected for stage-associated behavior and included 10 healthy controls and 43, 14, 13 and 6 stage I–IV HCC samples. Because the peak subset was selected on the same samples that then supply the trend, this panel is reported as exploratory and does not constitute an independent test of a stage association [82]; its overlap densities are on a different scale from the whole-peak-set analysis and the two are not directly comparable.

For tissue–plasma concordance we compared CpG-island methylation changes in HCC tumor versus adjacent non-tumor tissue with plasma-versus-healthy-control changes, computed before and after read selection. Pearson correlations were tested against zero, and directional quadrants were defined by the signs of the tissue and plasma methylation changes. Because the two correlations share both the tissue contrast and the CpG-island units, each coefficient and its test against zero are reported without an inferential comparison between coefficients; the redistribution across quadrants is reported as the summary of what selection changed. Differential CpG sets defined independently from HCC versus adjacent tissue and from CDX versus white-blood-cell comparisons were then evaluated in the selected and background pools.

Reference-category analysis tested the top 250 nominated genes against cell-type reference gene sets and did not estimate cell fractions. Promoter-level overlap between genes nominated from selected plasma reads and from liver-cancer CDX material was assessed for significance with a hypergeometric test against the set of promoters with sufficient coverage in both datasets, since overlap counts are not interpretable without the size of the shared universe [56].

### 4.13 Stage III pan-cancer analyses

We applied the frozen scorer to 348 independent plasma samples, comprising 58 healthy controls and 290 stage III cancer samples across 11 cancer types. The cancer samples included breast (n = 28), colorectal (n = 54), liver (n = 23), lung (n = 30), non-Hodgkin lymphoma (n = 18) and ovarian cancer (n = 20). The remaining samples comprised pancreatic (n = 19), bladder (n = 23), prostate (n = 8), esophageal (n = 28) and gastric cancer (n = 39). Only stage III cancer samples were included in this analysis. Pre-existing cohort labels were used for cohort summaries and stratified downstream analyses, never to retrain or recalibrate the scorer.

All-read and selected-read CpG-island profiles were projected separately by tSNE for inspection, and the difference between them was quantified by clustering the same profiles in the full feature space and scoring the result against tumor-type labels with the adjusted Rand index. Analysis-specific aggregation defined the ctDNA-BERT-derived index; this quantity was not calibrated against any external clinical measurement, is not zero in healthy controls and is not comparable with the pathology-calibrated FFPE estimates. Group comparisons of the index against healthy controls used two-sided rank-based tests with Benjamini–Hochberg adjustment and are reported with effect sizes as well as P values. Selected and background reads were intersected with cancer-matched ATAC-seq peaks for eight cancer types, with overlap normalized per 10^4^ valid reads and per megabase of peak territory. Promoter-associated methylation differences were mapped to genes and annotated with external tumor–normal expression and survival data from the TCGA cohorts; those external statistics derive from TCGA tissue data and are independent of the plasma cohort composition.

### 4.14 Stage I case-control and tissue-of-origin modeling

The stage I analysis used 1,477 specimens comprising 747 healthy controls and 730 cancers, all clinically staged as stage I at enrollment: 60 breast, 93 colorectal, 75 liver, 286 lung and 216 thyroid. Lung and thyroid cancer together account for 502 of the 730 cases, so aggregate performance is weighted toward those two types. Binary and cancer-type analyses were performed at sample level, and all feature selection, scaling and model fitting were contained within the corresponding cross-validation training folds.

Figures 6c,d were generated from the same five-fold stratified out-of-fold analysis of the 1,477-specimen cohort. The input dictionary comprised 5,489 variables, consisting of 5,456 k-mer frequencies plus 33 BED-derived annotations. Within each training fold, ANOVA retained 2,000 variables before StandardScaler and a class-balanced radial-basis-function support-vector machine (C = 20, *γ* = 1/d, seed 42) [83]. The same pooled out-of-fold scores were used to construct the ROC curves in Fig. 6c and the sensitivity-specificity traces in Fig. 6d. Pooled out-of-fold thresholds were set immediately above the ordered healthy-control score at each displayed specificity, so those thresholds were chosen post hoc rather than locked in advance.

The out-of-fold AUROC, sensitivity and specificity describe discrimination in this retrospective case-control cohort. These results do not directly establish positive predictive value in a screening population, which also depends on cancer prevalence. Cancer-type modeling used the 730 cancer cases in a separate nested five-fold out-of-fold analysis. The 5,482-feature raw representation contains 5,456 k-mer frequencies plus 26 non-k-mer summaries: 18 fragment-length, 5 methylation and 3 sequence-composition features. Adding the 33 BED-derived annotations to the 5,456-k-mer subset produces the 5,489-input dictionary used in Fig. 6c,d; adding them to all 5,482 raw features produces 5,515 inputs. The inner loop compared these feature blocks and class-balanced classifiers using macro F1. Logistic regression with C = 0.3 was selected in every outer fold. Folds 1, 2, 4 and 5 used the 5,489-input k-mer-plus-BED set while fold 3 used the 5,482 raw features; this difference was an outcome of inner-loop model selection rather than a manual change to a held-out fold. Standardization and model fitting were confined to training data, and held-out class probabilities were pooled for the confusion matrix and one-versus-rest ROC curves.

### 4.15 Serial colorectal-cancer analysis

Seven patients with colorectal cancer contributed serial specimens at T-01, T-02 and T-04. Figure 6h uses the ctDNA-BERT-derived index for the seven observed T-01/T-04 pairs and a two-sided paired test; the number of patients in whom the index decreased is reported alongside the P value. T-02 and gene-level promoter methylation summaries are shown descriptively in Supplementary Fig. 5; individual-patient promoter coverage and methylation comparisons are shown in Supplementary Fig. 6. No threshold-defined clinical response endpoint or monitoring operating characteristic was evaluated, so the serial analysis is limited to within-patient visualization and an exploratory paired comparison.

### 4.16 External annotations and statistical analysis

External annotations were introduced after read scoring. Cancer-type ATAC-seq peaks were derived from Data S4 of Corces et al. [53]. Motif intervals used UCSC annotations and the JASPAR 2024 CORE vertebrate collection [52]. The mappability-matched audit used the UCSC/ENCODE CRG 100-mer hg19 track [84].

Verified TCGA promoter-direction analyses used HumanMethylation450 matrices for BRCA, COAD, LIHC, LUAD and THCA and the corresponding hg19 GPL16304 probe map from the UCSC Xena TCGA Hub. The LIHC resource corresponds to the TCGA hepatocellular-carcinoma cohort [85]. Resource access details are recorded in the companion dataset inventory. Tissue methylation profiles were obtained from cited public resources. For the cell-type category analysis in Fig. 4g, enrichment results were obtained using the Human Gene Atlas gene-set library via enrichR. Genomic intersections used BEDTools or equivalent interval operations [86].

Tests were two-sided unless stated otherwise. Rank-based tests were used for non-Gaussian independent distributions, paired rank-based tests for linked longitudinal samples and for within-read category comparisons, and Pearson correlation for tissue– plasma concordance. Multiple comparisons were adjusted by the Benjamini–Hochberg procedure. Figure 6b used one-sided Mann–Whitney U tests with Benjamini– Hochberg adjustment. Methylation-token and motif-token attention were analyzed independently, without a CpG–motif adjacency or overlap requirement. Box plots show the median, first and third quartiles, with whiskers extending to the most extreme value within 1.5 times the interquartile range unless a panel states otherwise. Gene symbols follow HGNC nomenclature.

## Supplementary information

Supplementary Figs. 1–6 and their legends accompany this manuscript as a separate PDF. They provide training and attention analyses, additional mixture benchmarks, HCC methylation profiles and cell-type annotations, promoter-candidate analyses, and longitudinal colorectal cancer measurements.

## Declarations

### Data availability

The data reported in this paper have been deposited in the OMIX, China National Center for Bioinformation / Beijing Institute of Genomics, Chinese Academy of Sciences (https://ngdc.cncb.ac.cn/omix: accession no. OMIX022164). The deposit covers the study-generated Primer-RRBS and WGBS libraries analysed in this work, spanning the pretraining, tissue fine-tuning, xenograft, FFPE, HCC, stage III pan-cancer, stage I case–control and serial cohorts (Table 1).

Each library is provided as a gzip-compressed mHapX file that records, for every read, its genomic coordinates, strand, CpG methylation pattern and associated sequence, including the 3^*′*^ terminal sequence used as model input. These files are sufficient to rerun ctDNA-BERT scoring and to reproduce the read-level and sample-level analyses reported here.

Because these data derive from human participants, they are released under controlled access in accordance with the Regulations of the People’s Republic of China on the Administration of Human Genetic Resources and the scope of participant consent. Raw sequencing files are not included in the deposit because their release falls outside the scope of the human genetic resources approval; the mHapX files nevertheless retain all read-level information required for the analyses reported here. Researchers may request access through the OMIX controlled-access system.

The OMIX platform and CNCB-NGDC database resources are described in the corresponding database publications [87, 88].

Source data for Figs. 1–6 and Supplementary Figs. 1–6 are provided with this paper. De-identified cohort manifests, the composition and sampling weights of every in silico mixture, sample-level ctDNA-BERT predictions and the statistical source tables underlying each main and supplementary panel are provided as Supplementary Data.

Publicly reused RRBS reference data are available from NCBI GEO under GSE105127, GSE114254, GSE114807, GSE118255, GSE130582, GSE135770, GSE151527, GSE160331, GSE164365, GSE165360, GSE59163, GSE66167, GSE71702 and GSE93383; the 177 GSM-level records used are enumerated in Supplementary Data 1. Cancer-type ATAC annotations are available in Data S4 of Corces et al. JASPAR 2024 CORE vertebrate motifs were obtained from JASPAR. TCGA Human-Methylation450 matrices and the GPL16304 hg19 probe map were obtained from the UCSC Xena TCGA Hub. The UCSC/ENCODE CRG 100-mer hg19 mappability track was used for the mappability audit. These public resources did not alter the read-level ctDNA-BERT checkpoint; their analysis-specific downstream uses are described above.

## Code availability

The source code for ctDNA-BERT is hosted on GitHub at https://github.com/zxc-CCC/ctDNA-BERT.

## Author contributions

X.Z., X.S., R.W., N.Z. and F.L. contributed equally to this work. F.Z. and T.W. conceived and initiated the study. F.Z., T.W., X.Z. and X.S. designed the model and analyses. J.S. designed the analyses for model application. X.Z. implemented the model and performed modelling analyses. X.Z., X.S. and R.W. analysed the data and generated the results. R.W. and J.S. developed the mHapX software and curated the data. F.L. and N.Z. prepared the CDX and PDX samples and sequencing data. H.G. provided the experimental platforms and resources for the CDX and PDX studies. X.L. contributed to data analysis and figure preparation. J.H., Jiamin Sun, H.L., S.L., H.W., H.Z., H.S. and P.Z. contributed to modelling, data quality control, analysis and framework discussions. X.Z., J.S. and F.Z. wrote the manuscript. H.G., F.Z., J.S. and T.W. supervised and funded the work. All authors discussed the results and reviewed and approved the manuscript.

