## Supplemental figures for "Xenograft supervision and joint methylation-fragment-end encoding define a reusable molecular scoring layer for cell-free tumor DNA"

### Supplementary Fig. 1

Training, ablations and complementary attention examples.

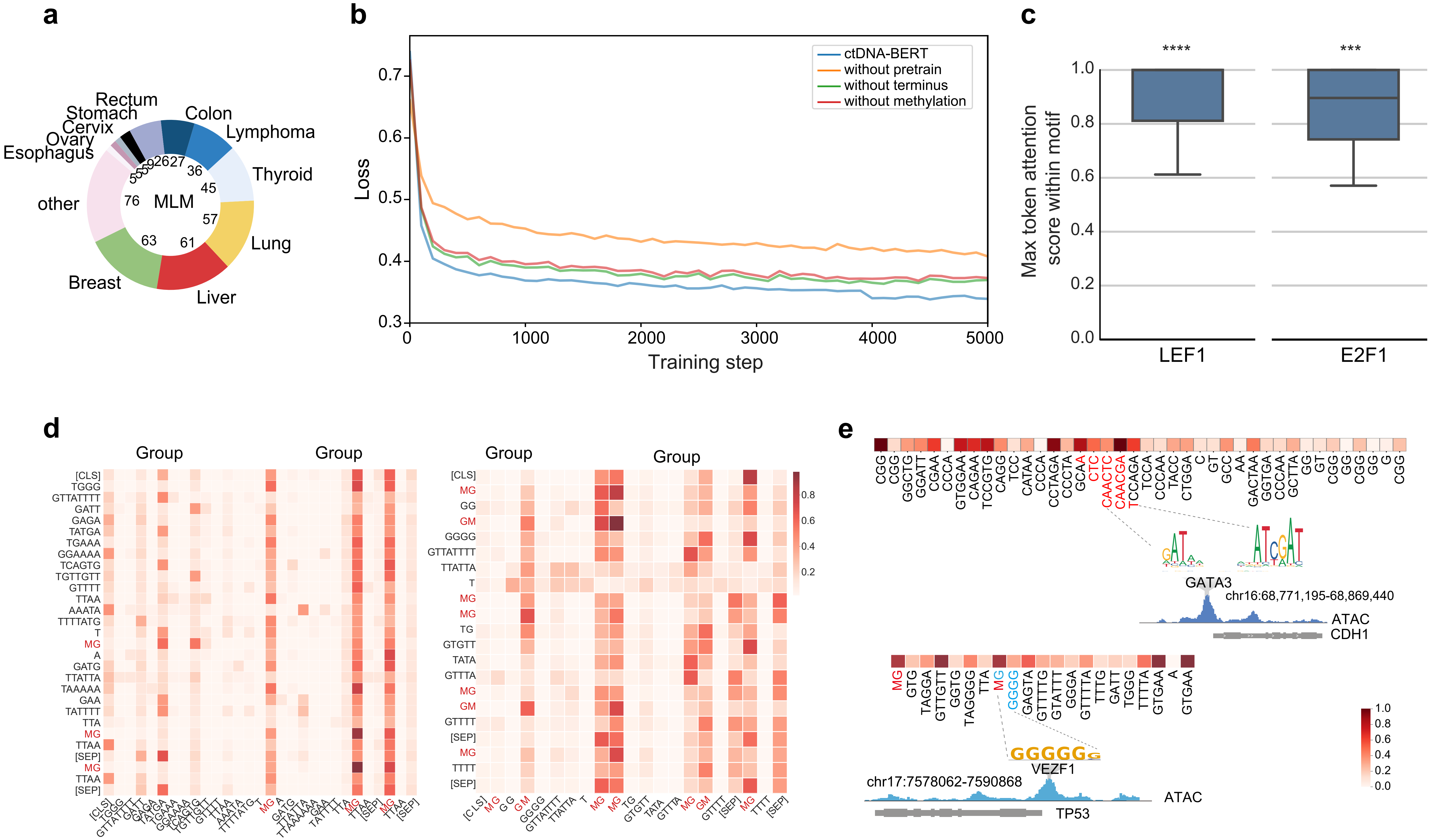


**Supplementary Fig. 1 | Training, ablations and complementary attention examples.** a, Category composition of the 415 libraries used for masked-language pretraining. b, Training-loss trajectories forcomplete ctDNA-BERT and variants without pretraining, the terminal segment or methylation-aware encoding over 0-5,000 steps. c, Maximum within-motif token attention for LEF1 (n = 30 motif intervals) andE2F1 (n = 25) in eligible LIHC reads. Original significance markers refer to comparisons with the CTCF reference using two-sided Mann-Whitney U tests with Benjamini-Hochberg adjustment, as described in themanuscript. d, Token-to-token attention matrices for two example reads, including MG/GM tokens. e, Additional CDH1/GATA3 read-level attention example with ATAC-seq and H3K27ac annotations. Theseviews complement Figs. 1 and 2. Attention weights describe model input weighting and do not establish transcription-factor occupancy. MLM, masked language modelling.

### Supplementary Fig. 2

Fraction recovery and mixture-dependent performance in complementary benchmarks.

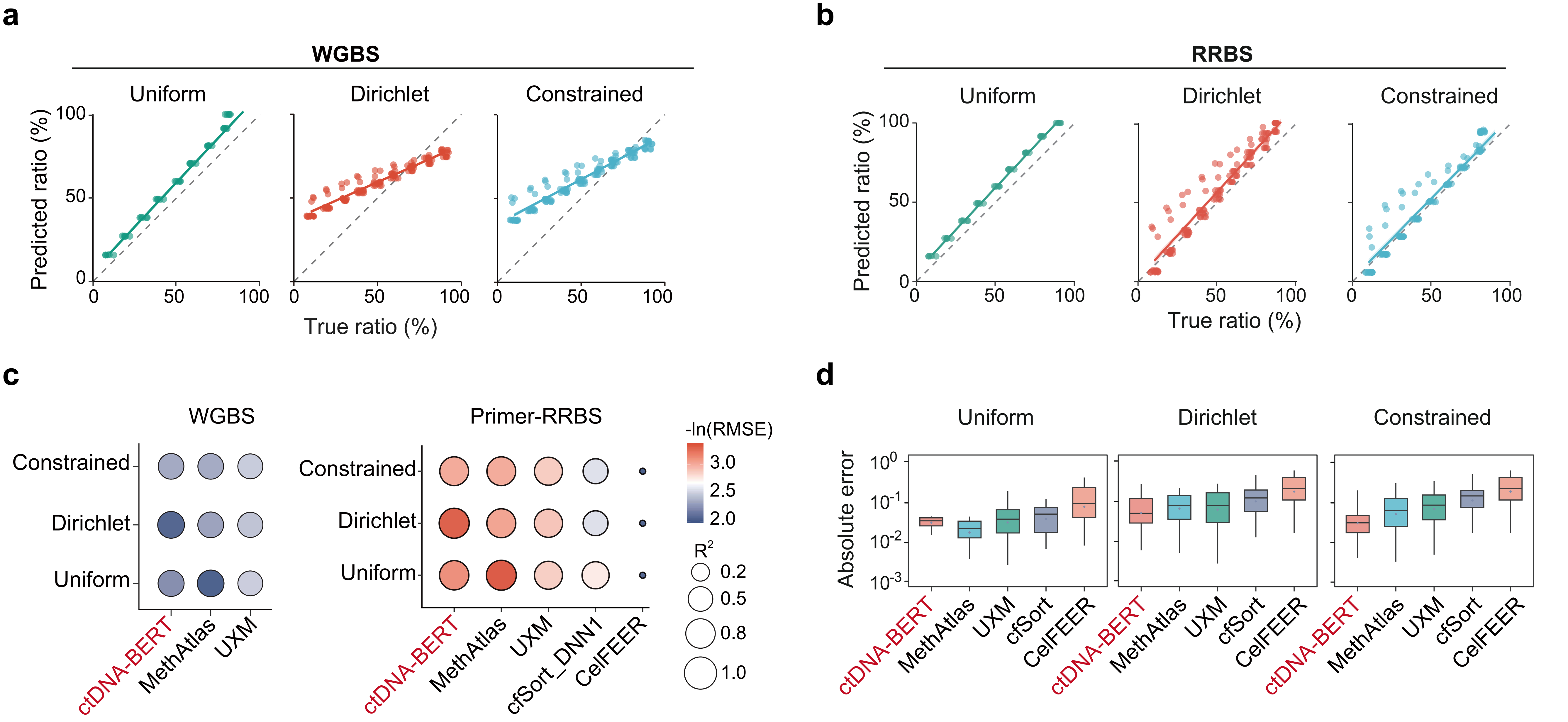


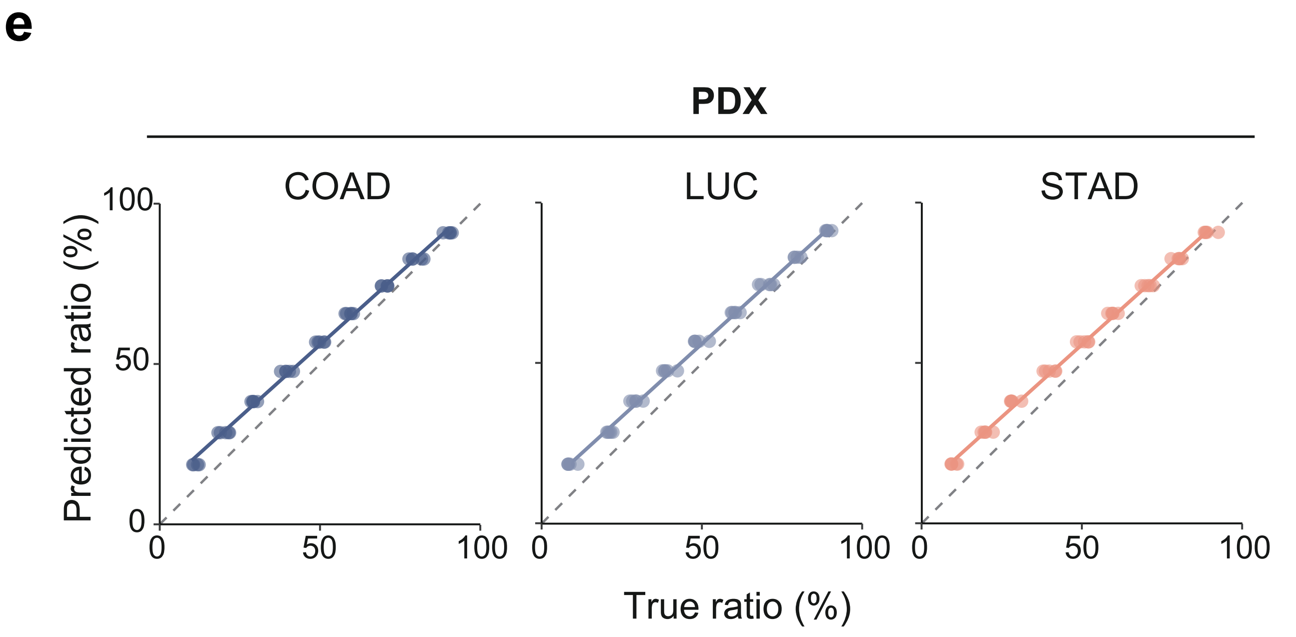


**Supplementary Fig. 2 | Fraction recovery and mixture-dependent performance incomplementary benchmarks.** a,b, Predicted versus true tumour fractions in WGBS (a) andPrimer-RRBS (b) mixtures under Uniform, Dirichlet and Constrained source-compositionregimes. c, WGBS (left) and Primer-RRBS (right) performance by regime. The WGBScomparison includes ctDNA-BERT, methAtlas and UXM; the Primer-RRBS comparison-additionally includes cfsort_DNN1 and CelFEER, retaining the source display labels. Bubble arearepresents R-squared and colour represents -ln(RMSE), following the original shared key. d,Primer-RRBS absolute-error distributions for ctDNA-BERT, methAtlas, UXM, cfSort andCelFEER, separated by regime. e, Fraction recovery in held-out colorectal (COAD), lung (LUC)and gastric (STAD) PDX mixtures with white-blood-cell background. Dashed diagonals in a,b,eindicate equality. The RRBS and PDX designs each comprise nine fractions (0.1-0.9) and tenreplicates per regime or tumour type, respectively. These views complement Fig. 3a,b. WGBS,whole-genome bisulfite sequencing; RRBS, reduced-representation bisulfite sequencing; PDX,patient-derived xenograft; RMSE, root-mean-square error.

### Supplementary Fig. 3

Individual HCC methylation profiles and expanded gene-reference annotations.

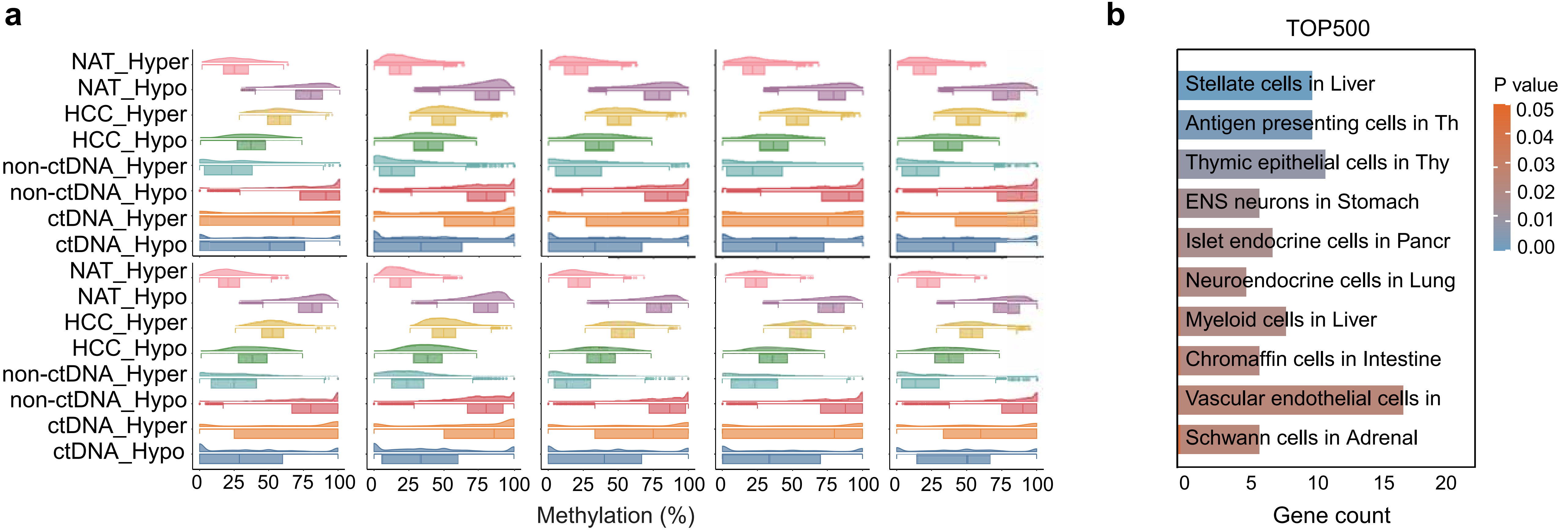


**Supplementary Fig. 3 | Individual HCC methylation profiles and expanded gene-reference annotations.** a, Differential-CpG methylation distributions in ten individual HCC plasma sample displays,retaining their original order. Reference hyper- and hypomethylated CpG sets were defined from HCC tissue versus non-tumour adjacent tissue (NAT), and from liver-cancer CDX reads versus white-blood-cell(WBC) background. Density curves and embedded box summaries show these sets in ctDNA-BERT-selected and background plasma reads, complementing the aggregate comparison in Fig. 4h. b, Reference-cell-type categories for the TOP500 HCC gene-candidate set. Bars show gene counts; colours follow the original P-value key. This expanded candidate-set annotation complements the TOP250 view in Fig. 4g.HCC, hepatocellular carcinoma; CDX, cell line-derived xenograft.

### Supplementary Fig. 4

Promoter methylation candidates and external bladder-cancer associations.

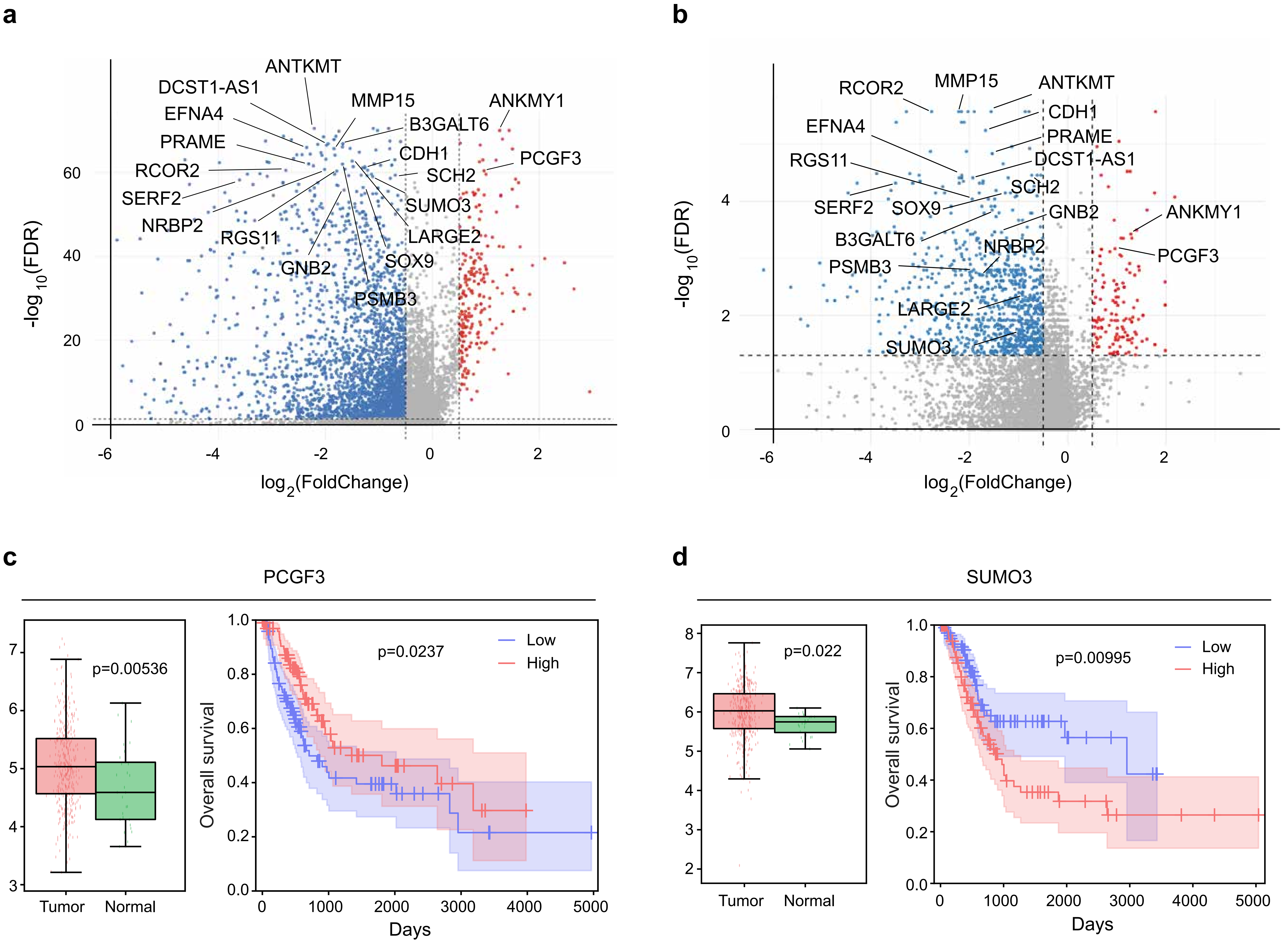


**Supplementary Fig. 4 | Promoter methylation candidates and external bladder-cancer associations.** Fig. 5e. b, Promoter-associated methylation differences between ctDNA-BERT-selected and background reads in the bladder-cancer analysis. Volcano plots retain the source fold-change and FDR axes and-candidate labels. c,d, External TCGA-BLCA tumour-versus-normal expression comparisons and overall-survival curves for PCGF3 (c) and SUMO3 (d). Survival curves separate high- and low-expressiongroups. The displayed expression and survival P values are 0.00536 and 0.0237 for PCGF3, and 0.022 and 0.00995 for SUMO3, respectively. The corresponding B3GALT6/SOX9 examples appear in Fig. 5f.FDR, false-discovery rate; BLCA, bladder urothelial carcinoma.

#### Supplementary Fig. 5

Serial sampling and gene-level promoter methylation in colorectal cancer.

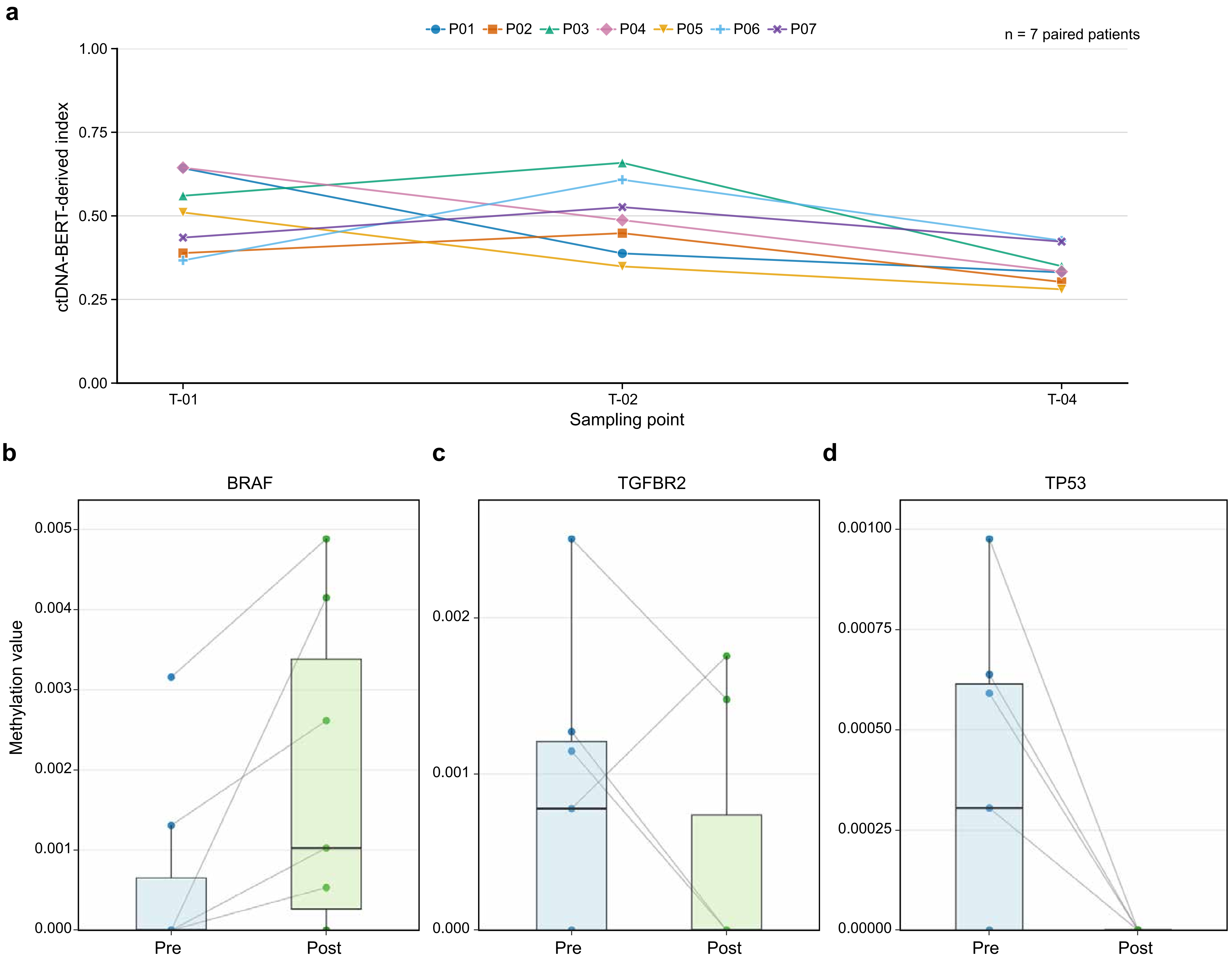


**Supplementary Fig. 5 | Serial sampling and gene-level promoter methylation in colorectal cancer.** a, ctDNA-BERT-derived index at T-01, T-02 and T-04 in seven colorectal-cancer patients; lines connectobservations from the same patient. The 21 observations provide the three-time-point context for the T-01/T-04 paired comparison in Fig. 6h. b-d, Paired pre-/post-treatment promoter methylation summaries forBRAF (b), TGFBR2 (c) and TP53 (d). P01-P07 identify the same patients throughout this figure and Supplementary Fig. 6. The index is the analysis-specific model-derived quantity reported in the manu-script.Individual-patient promoter coverage and methylation comparisons appear in Supplementary Fig. 6.

### Supplementary Fig. 6

Patient-matched promoter coverage and methylation before and after treatment.

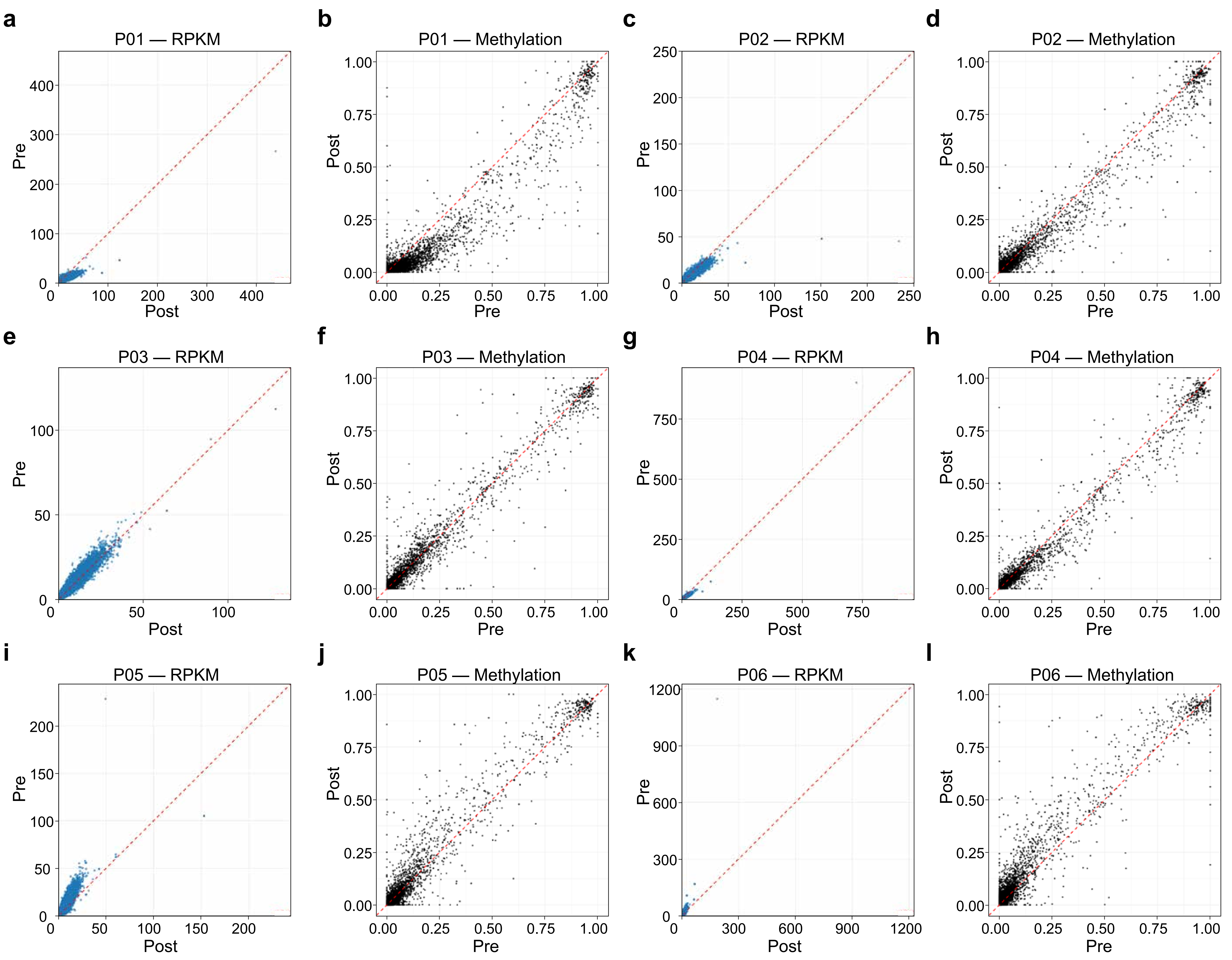


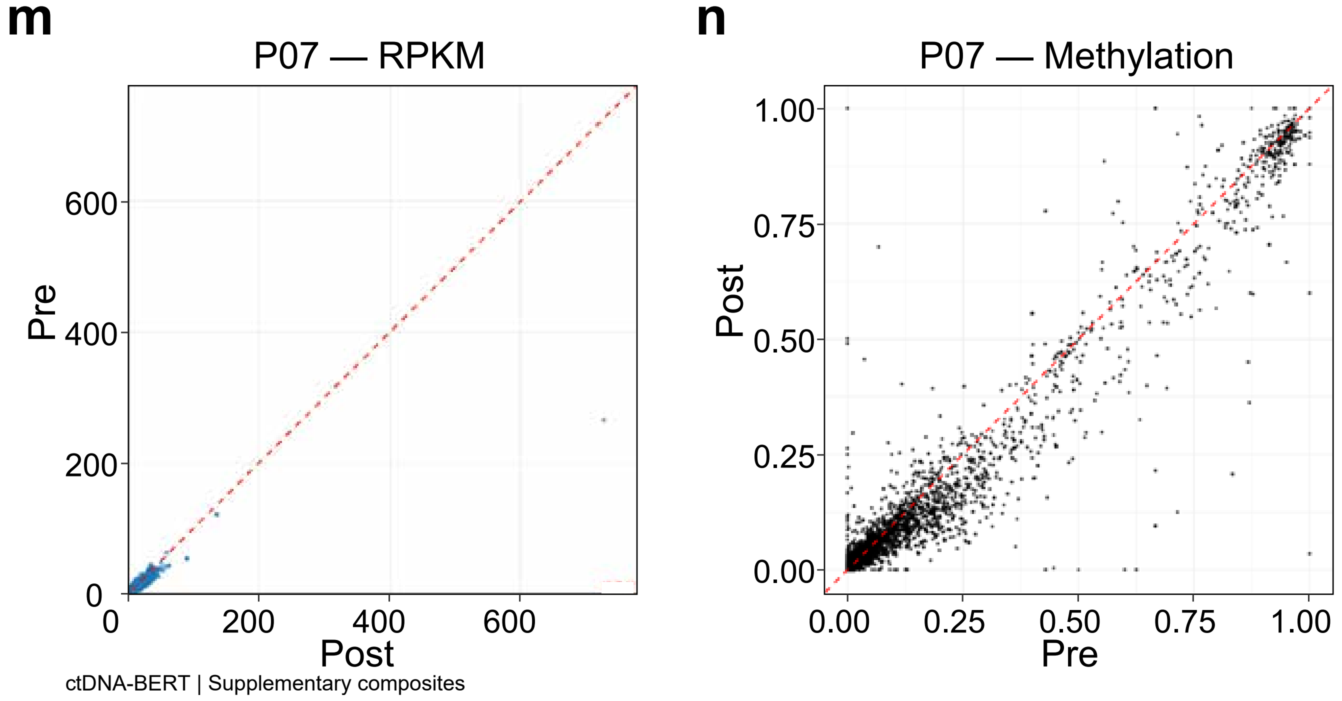


**Supplementary Fig. 6 | Patient-matched promoter coverage and methylation before andafter treatment.** a-n, Paired comparisons for colorectal-cancer patients P01-P07, with twoadjacent panels per patient. Coverage is shown in a,c,e,g,i,k,m; methylation is shown inb,d,f,h,j,l,n. Each point denotes a promoter-associated region. Coverage panels showpost-treatment average RPKM on the horizontal axis and pre-treatment average RPKM on thevertical axis. Methylation panels show pre-treatment methylation on the horizontal axis andpost-treatment methylation on the vertical axis. Red dashed lines indicate equality. Originalpatient-specific scales are retained. These descriptive within-patient comparisons complement Fig.6h and Supplementary Fig. 5. RPKM, reads per kilobase per million mapped reads.
